# Genetic signatures of divergent selection in European beech (*Fagus sylvatica L.*) are associated with the variation in temperature and precipitation across its distribution range

**DOI:** 10.1101/2021.05.18.444720

**Authors:** D. Postolache, S. Oddou-Muratorio, E. Vajana, F. Bagnoli, E. Guichoux, A. Hampe, G. Le Provost, I. Lesur, F. Popescu, I. Scotti, A. Piotti, G.G. Vendramin

## Abstract

High genetic variation and extensive gene flow may help forest trees with adapting to ongoing climate change, yet the genetic bases underlying their adaptive potential remain largely unknown. We investigated range-wide patterns of potentially adaptive genetic variation in 64 populations of European beech (*Fagus sylvatica* L.) using 270 SNPs from 139 candidate genes involved either in phenology or in stress responses. We inferred neutral genetic structure and processes (drift and gene flow) and performed differentiation outlier analyses and gene-environment association (GEA) analyses to detect signatures of divergent selection.

Beech range-wide genetic structure was consistent with the species’ previously identified postglacial expansion scenario and recolonization routes. Populations showed high diversity and low differentiation along the major expansion routes. A total of 52 loci were found to be putatively under selection and 15 of them turned up in multiple GEA analyses. Temperature and precipitation related variables were equally represented in significant genotype-climate associations. Signatures of divergent selection were detected in the same proportion for stress response and phenology-related genes. The range-wide adaptive genetic structure of beech appears highly integrated, suggesting a balanced contribution of phenology and stress-related genes to local adaptation, and of temperature and precipitation regimes to genetic clines. Our results imply a best-case scenario for the maintenance of high genetic diversity during range shifts in beech (and putatively other forest trees) with a combination of gene flow maintaining within-population neutral diversity and selection maintaining between-population adaptive differentiation.

## Introduction

Local adaptation is pervasive in forest tree populations (Alberto, Aitken, Alía, González-Martínez, et al., 2013; Savolainen, Pyhäjärvi, & Knürr, 2007) and is expected to play a major role in their response to ongoing environmental changes (Fady et al., 2016). Local adaptation implies that some key adaptive traits are genetically differentiated among populations, and thus that individual populations could evolve differently to the same environmental stress due to their different genetic setup (Kawecki & Ebert, 2004). Most temperate tree species have developed their present-day geographical patterns of local adaptation following considerable a range expansion from their glacial refugia after the Last Glacial Maximum (19.5-26 kyr BP, Clark et al., 2009; de Lafontaine, Napier, Petit, & Hu, 2018). There is ample concern that population processes going along with this postglacial range expansion such as founder events, genetic drift, or allele surfing might have left lasting imprints that could compromise the correct identification of adaptive genetic variation in extant natural tree populations (de Villemereuil, Frichot, Bazin, François, & Gaggiotti, 2014; Hoban et al., 2016; Rellstab, Gugerli, Eckert, Hancock, & Holderegger, 2015). Yet very few surveys of adaptive genetic variation in forest trees have to date assembled two prerequisites to properly account for species’ postglacial population dynamics: (i) a rangewide perspective with a sampling that thoroughly replicates populations from different regions and (ii) independent palaeoecological evidence documenting the species’ postglacial range dynamics and expansion (de Lafontaine et al., 2018). Empirical research to elucidate the issue is urgently needed because knowledge on local adaptation is crucial for conceiving conservation and management practices to adapt forest tree species and their ecosystems to ongoing environmental change (Aitken & Whitlock, 2013; Fady et al., 2016; Oney, Reineking, O’Neill, & Kreyling, 2013).

Reciprocal transplant experiments have been the classic approach to investigate local adaptation in forest trees, highlighting that phenotypes generally match their environment in extant populations (Rehfeldt et al., 2002; Savolainen et al., 2007). Provenance trials have also shown that trees generally have high levels of phenotypic plasticity at adaptive traits, and high levels of genetic variability within populations (Alberto et al., 2013; Gárate-Escamilla et al., 2019). This combination of local adaptation (i.e., mean trait values close to the optimum) and high within-population variation at key adaptive traits (i.e., large variance around the means) indicates a high genetic load, which in turn can be an asset when facing a swift environmental change (Savolainen et al., 2007). The development of population genomics has provided complementary approaches to study local adaptation (Hoban et al., 2016; Lind, Menon, Bolte, Faske, & Eckert, 2018). Two approaches, in particular, have become widely used to identify loci involved in local adaptation for non model organisms: differentiation outlier analyses, which aim at identifying loci with disproportionate allele frequency differentiation among populations, and gene-environment association (GEA) analyses, which aim at identifying loci exhibiting significant correlations with ecological variables. A key strength of such approaches is that they are cost-efficient for targeting large numbers of populations across well-defined environmental gradients, and therefore for nvestigating ecological hypotheses on the genomic basis of local adaptation (Capblancq et al., 2020; Rellstab et al., 2016; Temunović et al., 2020).

Present-day patterns of adaptive phenotypic and genetic differentiation have built up relatively quickly. It is only (at most) a few hundreds of generations ago that most temperate forest tree species recolonized large areas becoming available after the Last Glacial Maximum. Studies combining extensive surveys of fossil records (pollen and macro-remains) and of population genetic variation have been able to provide detailed direct evidence of the number and spatial location of glacial refugia, postglacial expansion routes, hybrid zones, and the timing of expansion events (e.g., de Lafontaine, Amasifuen Guerra, Ducousso, & Petit, 2014; Magri et al., 2006). This knowledge represents a highly valuable baseline information for disentangling demographic effects from those of selection, which is a major challenge for differentiation outlier and GEA analyses (de Villemereuil et al., 2014;Frichot, Schoville, de Villemereuil, Gaggiotti, & François, 2015; Hoban et al., 2016; Rellstab et al., 2015). Theoretical studies have shown that population expansions into new areas can go along with repeated founder effects, increasing random fluctuations of allele frequencies, possibly leading to the rise of neutral mutations to high frequencies (“allele surfing”), loss of genetic diversity and strong spatial genetic structure (SGS) along the expansion axis (de Lafontaine, Ducousso, Lefèvre, Magnanou, & Petit, 2013; Excoffier, Foll, & Petit, 2009; Slatkin, 1993). Allele surfing, in particular, could be mistaken for the increase in allelic frequency of a beneficial mutation propagated by selection (Paulose & Hallatschek, 2020; Ruiz Daniels et al., 2018). As both allele surfing and selection typically affect only a subset of loci in the genome, the former must be carefully considered when screening for the latter in expanding populations. Alternatively, colonization by many individuals should result in high genetic diversity at the colonization front and shallow SGS, particularly when founders originate from a variety of source populations. The effective number of founders depends on patterns of long-distance dispersal and on a variety of demographic processes and life-history traits (Austerlitz, Mariette, Machon, Gouyon, & Godelle, 2000; Fayard, Klein, & Lefèvre, 2009; Roques, Garnier, Hamel, & Klein, 2012). The actual relevance of these processes during the postglacial range expansion of temperate forest trees and their eventual traces in the present-day population genetic structures remain however under investigated.

This study takes advantage of the outstandingly well known postglacial population history of the European beech (*Fagus sylvatica* L.) to investigate the climate-associated genetic variation across its distribution range. Beech putative glacial refugia and colonization routes have been identified based on very detailed pollen records and genetic population surveys with chloroplast and isozyme markers (Magri et al., 2006). Beech is known to be highly sensitive to summer droughts (Aranda et al., 2015; Knutzen, Dulamsuren, Meier, & Leuschner, 2017), and, to a lesser extent, to late frosts (Kreyling et al., 2014; Petit-Cailleux et al., 2020). Genetic variation has been investigated at various climate-related phenological traits (Gárate-Escamilla, Hampe, Vizcaíno-Palomar, Robson, & Benito Garzón, 2019; Gauzere, Klein, Brendel, Davi, & Oddou-Muratorio, 2020; Gömöry & Paule, 2011; Kramer et al., 2017; Vitasse, Delzon, Bresson, Michalet, & Kremer, 2009), physiological or morphological traits (Bresson, Vitasse, Kremer, & Delzon, 2011; Hajek, Kurjak, von Wühlisch, Delzon, & Schuldt, 2016; Wortemann et al., 2011) and performance traits (Gárate-Escamilla et al., 2019). Phenological traits such as budburst and leaf senescence show consistent patterns of genetic variation across latitude or elevation at various spatial scales, with populations of higher elevation or latitude flushing earlier than populations from low elevation or latitude in common garden conditions (Gauzere et al., 2020; Gömöry & Paule, 2011; Vitasse et al., 2009). Genetic variation for performance traits, such as growth and juvenile survival, also shows a spatial structure, driven by spatial variations of maximal potential evapotranspiration (Gárate-Escamilla et al., 2019). By contrast, other functional traits involved in photosynthesis and transpiration are usually only weakly differentiated among populations but show instead high within-population variation (Hajek et al., 2016). These contrasting patterns of differentiation raise questions about the role of phenological and physiological traits in local adaptation and the spatial scales at which it takes place. Only a few published studies have so far used genetic approaches to investigate local adaptation, and mostly at local to regional scales (Capblancq et al., 2020; Csilléry et al., 2014; Cuervo-Alarcon et al., 2021; Krajmerová et al., 2017; Lalagüe et al., 2014; Müller, Seifert, & Finkeldey, 2015; Pluess et al., 2016).

This study investigates range-wide patterns of adaptive genetic variation in beech using 405 SNPs that are located in candidate genes involved either in budburst phenology and dormancy regulation (Lalagüe et al. 2014; Lesur et al. 2015) or in response to stresses (Lalagüe et al. 2014). We genotyped 446 individuals from 64 populations covering the entire species range (Fig. 1) to address the three following questions: (Q1) What are the risks that past population demography, including post-glacial recolonization, blur potential selective imprints on genetic structure? We expect limited genetic drift and allele surfing in beech. (Q2) Do certain loci show imprints of local adaptation? We expect genes related to phenological traits to show stronger signals of selection compared to genes related to stress-response traits. (Q3) if the spatial and climatic effects can be separated, what are the respective impacts of temperature- versus precipitation-related variables on adaptive differentiation? In line with Q2, we expected the temperature variables to stand out more than precipitation variables.

**Figure 1:**
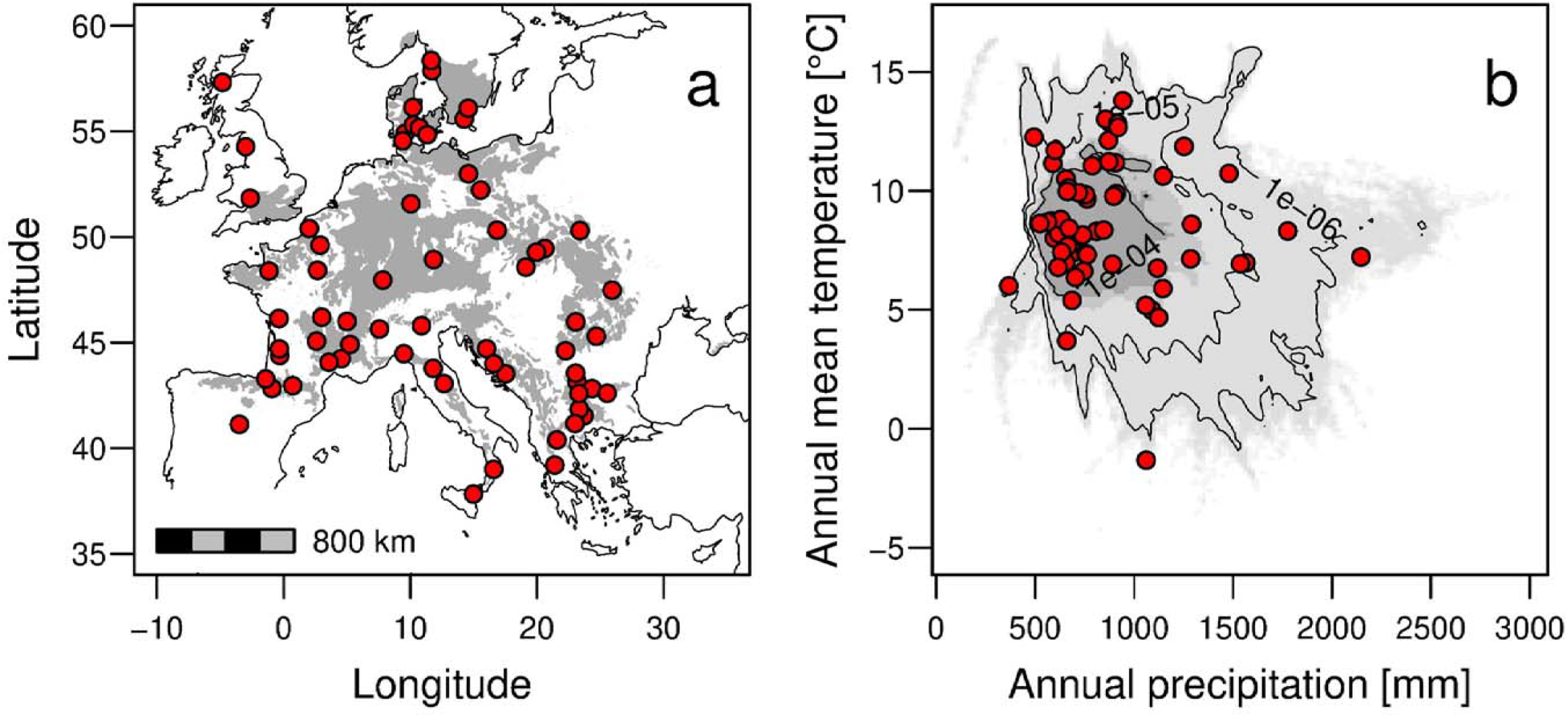
Distribution of the 64 studied populations (red dots) (a) in the geographical space, overlaid on beech distribution range (in grey) ; (b) in the bioclimatic niche defined by annual precipitations and temperature, with the grey colour intensity indicating increasing density of beech stands.

## Materials and Methods

### Sampling design

European beech is a dominant broadleaved tree species of many lowland and mountain forests across Europe, extending from Spain to the Carpathians and from Sicily to southern Sweden. We sampled 446 adult trees in 64 populations across Europe (Fig. 1A), thoroughly covering the geographical range and bioclimatic niche of beech (Fig. 1B) and including all the major glacial refugia identified by Magri et al. (2006). Leaves were collected from 4 to 10 (7 on average) haphazardly chosen dominant adult trees (at a minimal distance of 40 m from each other) growing in native beech stands.

### SNP development, genotyping and filtering

Nuclear DNA was extracted from 20-30 mg of dry leaf tissue per individual with the DNeasy Plant Mini Kit (QIAGEN) following the manufacturer’s instructions. DNA concentration was measured on a ND-8000 NanoDrop spectrophotometer (Thermo Scientific, Wilmington, USA). Samples were genotyped at 405 SNPs distributed in two multiplex assays.

The first assay of 165 SNPs previously developed by Lalagüe et al. (2014) was carried out using Kompetitive Allele Specific PCR (KASP; He, Holme, & Anthony, 2014). Among those, 37 SNPs were located in 15 genes annotated as phenology-related in *Quercus petraea*. The other 128 SNPs were located in 37 stress-related genes that were selected from different sources (see Lalagüe et al., 2014 for details) : (1) literature on candidate genes involved in plant response to abiotic stress; (2) sequences of three proteins involved in cavitation resistance in beec); (3) annotated amplicons from *Q. petraea* and *Q. robur*.

The second assay targeted 240 SNPs in six multiplexes of 40 SNPs each. This assay was developed for this study from available genomic resources (Lesur et al., 2015) and included 104 SNPs located in 51 genes differentially expressed in quiescent buds (QB) as well as 116 SNPs located in 58 genes differentially expressed in swelling buds (SB). This assay also included 20 unrelated control SNPs located in 15 housekeeping genes. Genotyping was performed on a MassARRAY System (Agena Bioscience, USA) using the iPLEX Gold chemistry following Gabriel *et al*. (2009). Data analysis was performed with Typer Analyzer 4.0.26.75 (Agena Bioscience). We filtered out all monomorphic SNPs, as well as loci with a weak or ambiguous signal (i.e., displaying more than three clusters of genotypes or unclear cluster delimitation).

Raw variant data were filtered with *Plink* (v.1.9; Chang et al., 2015). Individuals and SNPs with >15% missing data were filtered out, together with variants with MAF<1%.

### Linkage disequilibrium

Pairwise linkage disequilibrium (LD) estimates were obtained within each of the genetic clusters identified by genetic clustering analyses (see below) using the *LD()* function in the R package *genetics* (Warnes, Gorjanc, Leisch, & Man, 2021). The *p*.*adjust()* function was used to correct *p*-values for multiple testing with the Bonferroni method, and association between allele frequencies was deemed as statistically significant at a nominal significance threshold equal to 1⍰10^−3^. An *ad hoc* algorithm was devised to iteratively identify and remove the loci most frequently involved in pairwise significant tests, leading to cluster-specific lists of SNPs showing statistical evidence of linkage. The markers shared across all analyses were removed to obtain an LD-pruned version of the SNP dataset to be used in the analyses assuming linkage equilibrium among loci (*i*.*e*., STRUCTURE and pcadapt, see below).

### Population genetic structure

The Bayesian clustering analysis implemented in STRUCTURE (Pritchard, Stephens, & Donnelly, 2000) and the multivariate method implemented in the Discriminant Analysis of Principal Components (DAPC; Jombart, Devillard, & Balloux, 2010) were used to infer patterns of population structuring and admixture among beech populations. The main difference between the two methods is that STRUCTURE builds genetic clusters so to minimise the overall departures from HWE, whereas DAPC is based on maximizing the differentiation between inferred genetic clusters while minimizing variation within them.

STRUCTURE 2.3.4 (Pritchard et al., 2000) was run using default settings and parameter values, assuming the admixture model, and the putative number of different genetic clusters (K) ranging from one to 10. Each run consisted of 5×10^4^ burn-in iterations and 1×10^5^ data collection iterations. Ten independent runs were performed for each value of K. The average likelihood and ΔK statistics described in Evanno *et al*. (2005) were calculated for each K and used to identify the most-likely K-value. For informative values of K, distinct runs were averaged using CLUMPAK (Kopelman, Mayzel, Jakobsson, Rosenberg, & Mayrose, 2015) to obtain the final estimates of the membership coefficients (*q*-values) at individual and population levels. To comply with model assumptions, STRUCTURE was run first with the complete SNPs dataset, and then with the LD-pruned version of the dataset (once LD estimated within each cluster).

DAPC is based on a discriminant analysis (DA) of genetic data preceded by a few analytical steps to meet its requirements, all implemented in the R package *adegenet* (Jombart, 2008). Since DA requires *a priori* definition of clusters, K-means clustering of principal components (PC) on individual allele frequencies was first used to identify both group priors and the most likely number of genetic clusters. K-means was run on 150 PCs with K ranging from one to 40, and the Bayesian Information Criterion (BIC) was used to assess the best supported K-value. Then, as DA requires the variables to be uncorrelated and fewer than the number of observations, we used a principal component analysis (PCA) and the randomization approach implemented in the *a*.*score()* function in *adegenet* to select the number of PCs optimizing the trade-off between power of discrimination and over-fitting. Finally, the DA was run on 23 PCs extracted from the original dataset.

### Basic diversity and differentiation statistics

We computed allelic richness (*A*_*r*_) and mean number of alleles per locus (*N*_*a*_) using the R package *diveRsity* (Keenan, McGinnity, Cross, Crozier, & Prodöhl, 2013); percentage of polymorphic loci using GenAlEx 6.5 (Peakall & Smouse, 2012); observed (*H*_*O*_) and expected (*H*_*E*_) heterozygosity, Wright’s inbreeding coefficient (*F*_*IS*_), and *β*_*WT*_ (a plot-specific index of genetic differentiation relative to the entire pool; Weir and Goudet 2017) using the R package *hierfstat* (Goudet, 2005). Parameters of genetic fixation (*G*_*ST*_; Nei, 1977) and differentiation (Jost’s *D*; Jost, 2008) among populations/clusters were also calculated with GenAlEx and their statistical significance assessed with 999 permutations.

### Isolation by distance and barriers to gene flow

We estimated spatial genetic structure (SGS) among populations and tested whether geographic distance significantly shaped the patterns of genetic differentiation (as estimated by *F*_*ST*_*/1-F*_*ST*_) using the software SpaGeDi 1.4c (Hardy and Vekemans 2002). To test for isolation by distance (IBD), the (*F*_*ST*_*/1-F*_*ST*_) values were regressed on ln(*d*_*ij*_), where *d*_*ij*_ is the spatial distance between populations *i* and *j*. Then, we tested the regression slope (null hypothesis: *blog*_*FST*_ = 0) using 5,000 permutations of genotypes over populations. These analyses were run both on the 64 populations and within each cluster identified with DAPC. For within-cluster analyses, we retained all the individuals successfully assigned to a given cluster (i.e., those having a *q*-value above the nominal threshold of 0.6).

Spatial variation in genetic diversity and gene flow rates were estimated using Estimated Effective Migration Surfaces (EEMS; Petkova et al., 2016). This method tests for regional departures from the IBD model: areas where the decay of genetic differences across geographical distance is higher than expected under an IBD model are considered as suggestive of barriers to gene flow. A user-selected number of demes determines the geographic grid size and possible migration paths between all populations, and the EEMS are calculated by adjusting the migration rates so that the genetic differences obtained under a stepping-stone model match as closely as possible the observed genetic differences. The estimates are subsequently interpolated over the geographic space to provide a surface of observed genetic dissimilarities. We ran the runeems_snps executable with 500,000 burn-in MCMC steps and 2⍰10^6^ subsequent iterations. To reduce the potential influence of grid size, we averaged the results over nine independent runs with different numbers of demes (800, 1200 and 1600 with three repetitions each) and combined the results across the three independent analyses. We assessed convergence of runs, plotted geographic distance and genetic dissimilarity across demes, and generated effective diversity (q) and effective migration rates (m) surfaces using the R package *reemsplots* (Petkova et al., 2016).

### Bioclimatic data

Each sampling site was characterized by a set of 19 bioclimatic variables extracted from the WorldClim database v.1.4 (Hijmans, Cameron, Parra, Jones, & Jarvis, 2005) with a grid cell resolution of 30-arc second (ca 1×1 km) using DIVA-GIS v.7.5. These bioclimatic variables represent annual trends (e.g., mean annual temperature, annual precipitation), seasonality (e.g., annual range in temperature and precipitation) and extreme or limiting climatic factors (e.g., temperature of the coldest and warmest month, and precipitation of the wettest and driest quarter). To calculate predictors in the following analyses, we used the mean values of these 19 bioclimatic variables (Table S1) over the period from 1950 to 2000.

To reduce the multidimensional bioclimatic data set to a few uncorrelated factors, we performed two PCAs using the R package *FactoMineR* (Lê, Josse, & Husson, 2008), one focusing on the temperature-related variables (BIO1 to 11), and the other focusing on the precipitation-related predictors (BIO12 to 19). The selected principal components of each PCA were used as individual climate variables in *lfmm* and *Samβada* analyses, and combined in matrices to represent the climatic structure in the variance partitioning analyses (see sections below).

### Detection of signatures of selection

To detect loci carrying putative signatures of divergent selection (i.e., outliers), we used two differentiation outlier search methods, *pcadapt* (Luu, Bazin, & Blum, 2017) and the *F*_*ST*_-based method by Martins et al. (2016) as implemented in *lea* (Frichot & François, 2015). Moreover, we used two genotype-environment associations analyses, *lfmm* (Frichot, Schoville, Bouchard, & François, 2013) and *Samβada* (Stucki et al., 2017); these approaches are detailed in Supplementary Online Appendix 2. Note that each method allows the correction of outlier detection for the confounding effects of population structure. For all four methods, *p*-values were corrected across multiple tests using the same local False Discovery Rate (FDR) algorithm.

Finally, we annotated the identified outliers. For the loci obtained from Lalagüe et al. (2014), we queried against The Arabidopsis Information Resource (TAIR 11) database using BlastX with an E-value cut-off of 10^−5^. For the loci obtained from Lesur et al. (2015), the functional annotation of gene sequences containing outlier SNP was also reported (from their Table S2).

### Isolation by distance and environment

To evaluate the respective importance of IBD versus isolation by environment (IBE), we used a variance partitioning approach (Legendre, Fortin, & Borcard, 2015). We partitioned the explanatory power (as expressed by the adjusted *R*^2^) of the climatic and spatial structures on the genetic structure. Genetic structure was obtained through a Principal Coordinate Analysis (PCA) on the matrix of pairwise population genetic distances as returned by GenAlEx (Peakall & Smouse, 2012). Principal coordinates explaining up to 80% of the total variance were included in the response data table. The spatial structure was modelled by distance-based Moran’s eigenvector maps (dbMEM; Dray et al., 2006), as suggested by Legendre et al. (2015), and estimated by the *mem()* function in the R package *adespatial* (Dray et al., 2018). We retained only the statistically significant eigenvectors modelling positive spatial autocorrelation and, therefore, describing global patterns *sensu* Jombart et al. (2008). The climatic structure was summarized by temperature- or precipitation-related synthetic climatic variables (see “Bioclimatic data” section above). We assessed the relative contribution of climatic and spatial structure in explaining the genetic structure of populations using the function *varpart()* of the R package *vegan* (Oksanen et al., 2019). Significance of the variance components was calculated through an ANOVA-like permutation test for redundancy analysis (RDA) and partial ReDundancy Analysis (pRDA) based on 10,000 permutations (Legendre & Legendre, 2012).

All statistical analyses were conducted using R 3.6.2 (R Core Team, 2019) unless otherwise indicated.

## Results

### SNP dataset

Of the 405 SNPs from the two multiplex arrays, 135 were discarded from the analyses: 9 failed to amplify in all samples; 50 were monomorphic; 75 were of poor quality (visual clustering inspection); and one had a call rate <85%. Sixteen individuals were discarded due to a low call rate. The resulting dataset comprised 430 individuals and 270 SNPs (Supplementary Online Appendix A1), with 3-10 individuals per population (6.7 on average). The 270 SNPs included 150 SNPs in 93 phenology genes, 109 SNPs in 38 stress-related genes, and 11 SNPs in 8 housekeeping genes. After removing linked markers, the LD-pruned version of the dataset comprised 212 SNPs.

### Population genetic structure

Both the STRUCTURE and the DAPC analysis revealed an optimal grouping at K=3 (Fig. 2 and Fig. S1) with clear geographic boundaries among the three genetic clusters (as represented by the green, red and blue colors in Fig. 2) and admixture zones (areas with cross symbols in Fig. 2). Populations were considered as admixed when none of the population *q*-values exceeded the threshold of 0.60 for any inferred genetic cluster (Table S2).

**Figure 2:**
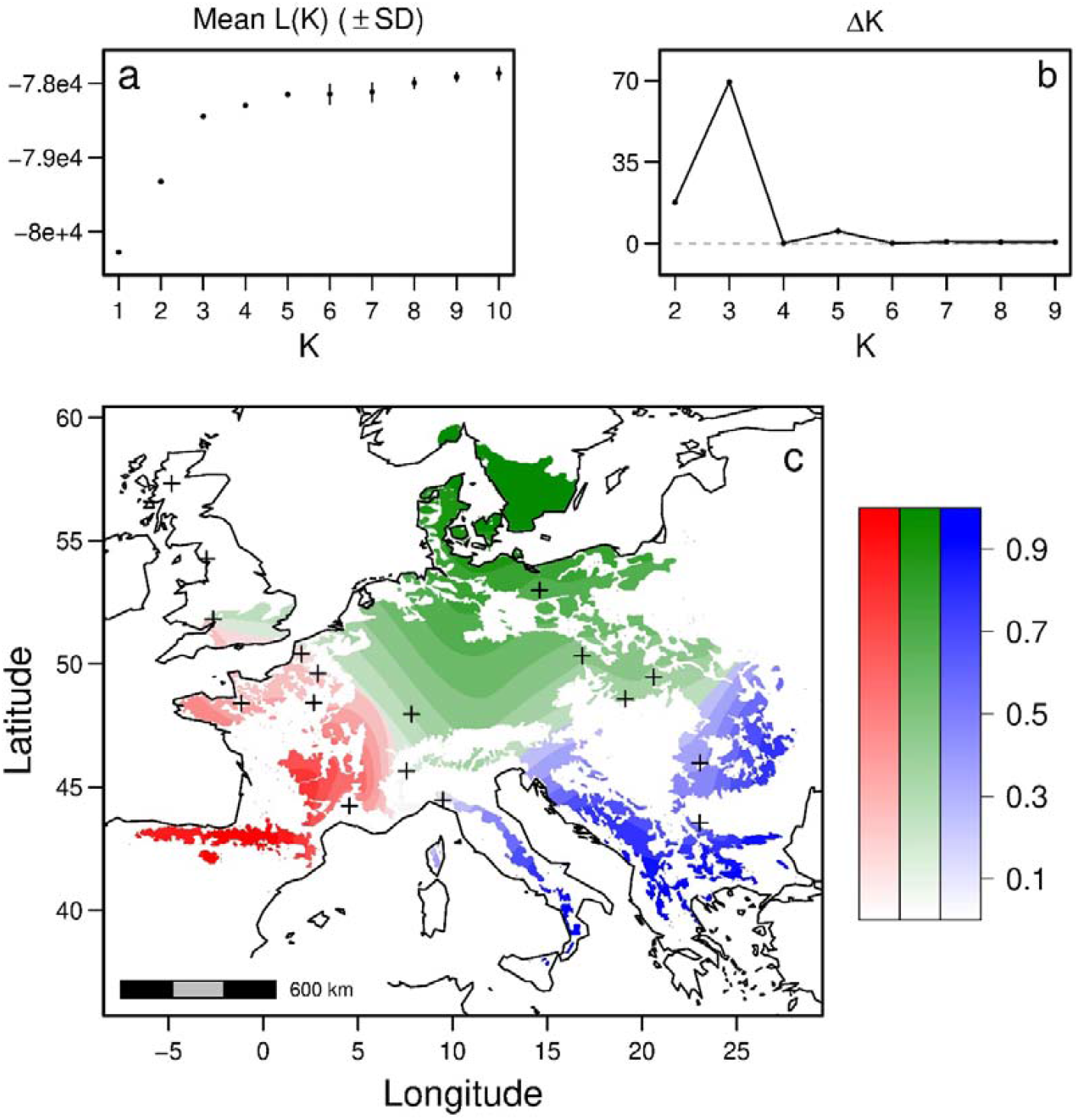
Spatial interpolates of the admixture coefficients estimated with STRUCTURE for K=3. Each color corresponds to one cluster and the colour intensity indicates the probability to belong to the cluster at a given position in space, based on spatial kriging of the individual q-matrix. Only areas belonging to beech distribution range are considered. Crosses indicate admixed populations, not assigned to a single cluster.

The green (dominant in Northern Europe) and red (dominant in Western Europe) clusters were separated by a main genetic boundary extending from the northern Tyrrhenian coast to southern England, which was also an area of admixture. In southern Europe, populations from the Apennines clustered with those from the Balkan peninsula and Carpathian mountains, forming the blue cluster. In central-eastern Europe, the main boundary between the blue and green clusters was located between Slovenia and Croatia and between the Western and Eastern Carpathian mountains. The area between the southern Carpathian mountains and the Baltic sea represented the second largest admixture area detected. The *G*_*ST*_ and Jost’s *D* pair-wise values among the three genetic clusters spanned from 0.024 to 0.025, and from 0.022 to 0.023, respectively.

### Spatial patterns of genetic diversity and differentiation

Genetic diversity and differentiation estimates are detailed in Table S2. The percentage of polymorphic SNPs within a population (%polloc) ranged from 62% to 87 % (mean %polloc = 77%), corresponding to an allelic richness ranging from 1.48 to 1.65 (mean Ar = 1.58). Observed and expected heterozygosities per population ranged, respectively, from 0.262 to 0.347 and from 0.248 to 0.327 (mean *H*_*o*_ = 0.305 and mean *H*_*e*_ = 0.300), indicating a small heterozygote excess (mean *F*_*IS*_ = -0.025). All genetic diversity indices showed the same patterns of variation across the range (Fig. 3 a, b), where southeastern populations had below-average values, whereas western populations showed above-average values. The indices of genetic diversity (*H*_*e*_, %polloc) also varied among clusters (Fig. S2). Populations assigned to red and green clusters had a higher *H*_*e*_ than populations assigned to the blue cluster (*p* = 0.003 and 0.074 respectively), while the red cluster populations showed higher %polloc values than those of the green and blue clusters (*p* < 10^−3^).

**Figure 3:**
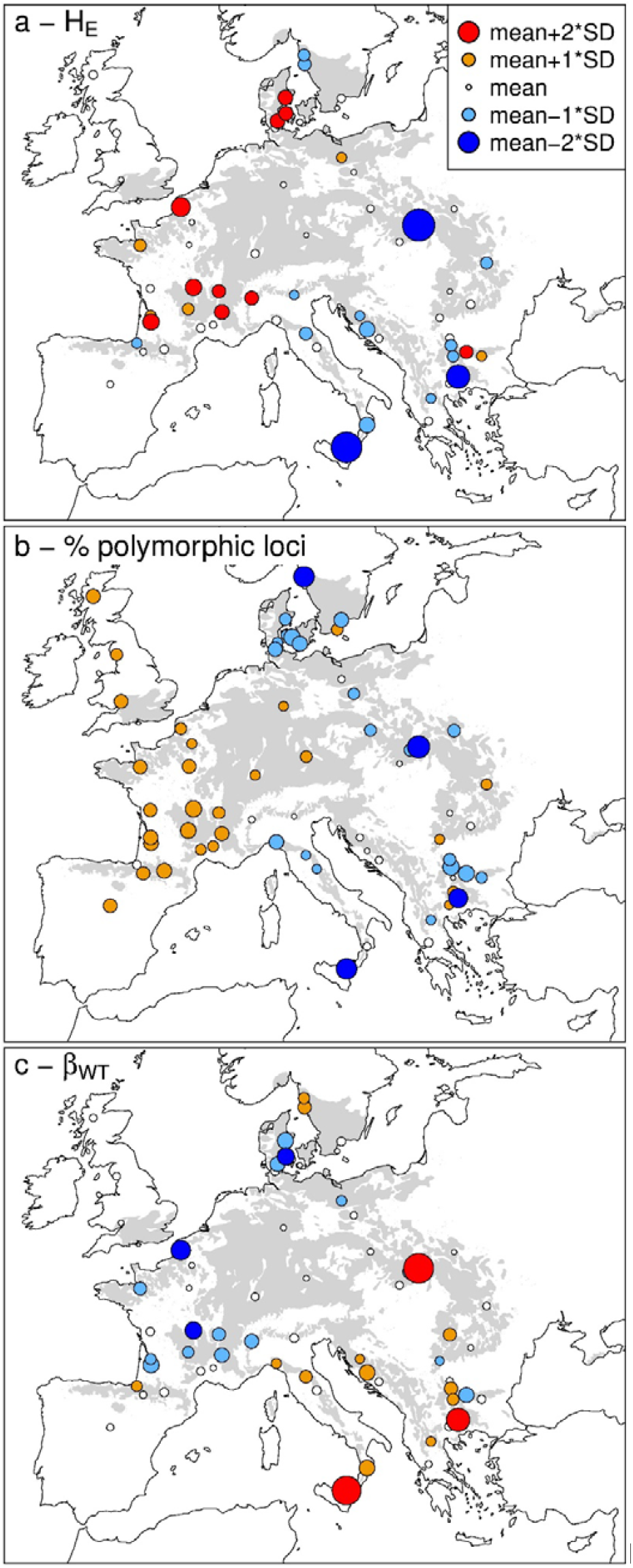
**Estimates of diversity (He), percentage of polymorphic loci (%polloc) and genetic differentiation relative to the entire pool (β_WT_) in the 64 studied populations, overlaid on beech distribution range (in grey)**.

Spatial patterns of genetic diversity were consistent when assessed independently with SNPs from the two different arrays (Fig. S3). Although other array-specific representation problems may occur, such a finding rules out a major distortion due to ascertainment bias.

Genetic differentiation of each population from the entire gene pool ranged from - 0.034 to 0.20 (mean *β*_*WT*_ = 0.05). Patterns of *β*_*WT*_ variation across Europe were opposed to diversity patterns (Fig. 3c), with southeastern populations characterized by higher average values than western populations. Accordingly, *β*_*WT*_ of the red and green cluster was higher than *β*_*WT*_ of the blue cluster (*p* = 0.005 and 0.091 respectively) (Fig. S2 c).

Genetic differentiation between all populations pairs revealed a significant signal of IBD (Fig. 4a). Pairwise *F*_*ST*_*/1-F*_*ST*_ values increased with increasing geographic distance (blog = 0.027, *p* < 0.001). This IBD signal was also observed within the blue and green clusters (Table S3, Fig. S4), but not within the red cluster.

**Figure 4:**
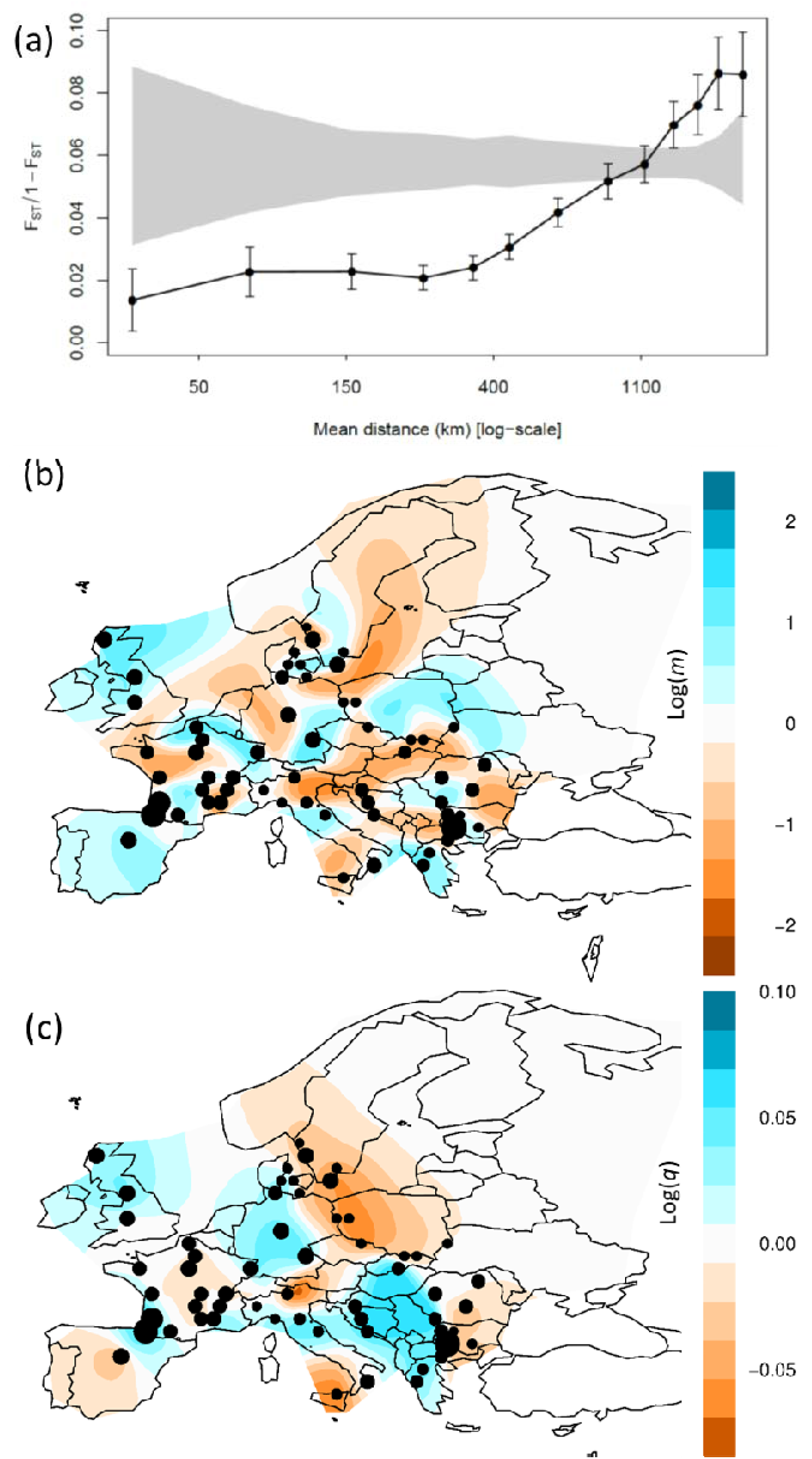
Patterns of isolation by distance and barriers to gene flow among the 64 studied populations. (a) Spatial genetic structure as depicted by the variation of genetic differentiation against geographic distance (on a log-scale). The grey envelope represents expected *F*_*ST*_/1-*F*_*ST*_ values under complete spatial randomness and bars represent standard error at 95% level within each distance class. (b) Contour maps representing the posterior mean of effective migration (*m*) surface; populations in the blue areas are connected by higher migration rates than expected under isolation by distance (IBD) while the ones in the orange areas have lower migration rates than expected and are interpreted as migration barriers. In white areas, the effective migration surface is close to the one expected under IBD. (c) Contour maps representing the posterior mean of effective diversity (*q*) surface; populations in the orange (respectively blue) areas have lower-than-expected (respectively higher-than-expected) genetic diversity than the average. On maps (b) and (c), black dots represent the studied populations, aggregated per grid cell (with size proportional to the number of genotyped individuals)

The EEMS analyses highlighted several barriers to gene flow corresponding to biogeographical barriers (Fig. 4b). The first barrier separated the UK and Scandinavia from the European mainland, and corresponded to the English Channel, the Baltic sea and plains in Western Germany and North-Western France. A second barrier corresponded to the Alps (Northern Italy, Austria) and extended to Slovakia and the Carpathians in the east. Weaker barriers to gene flow also occurred in Southern Italy and the Balkans.

Spatial patterns of effective diversity estimated with EEMS (Fig. 4c) partially contrasted with the maps of *H*_*e*_ and %polloc (Fig. 3). Indeed, areas of higher-than-average diversity were found with EEMS in Western Europe (UK, Pyrenees mountains, Germany) but also in Central and South-Eastern Europe. Diversity was lower than average in Spain, Southern Italy, and in an area from Eastern Scandinavia to Poland.

### Climate data analysis

The first three principal components of the temperature-focussed PCA (Temp1, Temp2, Temp3) were retained, and accounted for 90.9% of the total variance of the dataset (Online Appendix A3). Temp1 is an axis of mean temperatures, opposing hot (southern) to cold (northern) climates. Temp2 can be interpreted as an axis of climate continentality, opposing climates with strong versus weak variation of temperatures among years and seasons (e.g. continental *vs*. oceanic climates). Temp3 can be interpreted as an axis of climate xericity, opposing climates with a high diurnal range and the wettest season corresponding to the coldest months (i.e., mediterranean climates) to climates with a low diurnal range and the wettest season corresponding to the warmest months (i.e. temperate mesic climates).

The first three principal components of the precipitation-focussed PCA (Precip1, Precip2, Precip3) were also retained, and accounted for 98.6% of the total variance of the dataset. Precip1 is a precipitation abundance axis, opposing wet (Great-Britain, Northern Italy) to dry climates (Greece, Spain). Precip2 is a precipitation variability axis, opposing climates with strong (Greece, Italy) versus weak (France) variation of precipitation. Precip3 captures the coupling between precipitation and seasonal temperatures, opposing climate where high precipitation occurs during the vegetation period (Poland, Romania) to those where high precipitation occurs in winter (Greece, Italy).

### Selection signatures

#### pcadapt

The first two PCs were retained to represent population structure in *pcadapt* analysis based on the Cattle’s rule (Online Appendix A3). One candidate SNP under selection (0.3%) was identified after controlling for FDR (Table S4).

#### lea

The lowest cross-entropy criterion value was found at K = 3. Five SNPs (1.85%) were identified as potentially under divergent selection after controlling for FDR and after calibrating *p*-values using the calculated genomic inflation factor (λ = 6.0; Table S4).

#### lfmm

After controlling for FDR, 46 SNPs (17%) were found to be associated with temperature or precipitation-related climatic variables (60 significant associations in total, Table S5): seven SNPs showed correlations with Temp1, 13 with Temp2 and 11 SNPs with Temp3 ; four SNPs showed correlations with Precip1, nine with Precip2, and 16 with Precip3.

#### Samβada

Thirty-two genotypes at 22 SNPs (8.1%) were associated with temperature and precipitation-related variables after controlling for FDR. In particular, four loci showed correlations with Precip1 and 11 with Precip3, while 5 SNPs were associated withTemp1, seven with Temp2 and five with Temp3 (Table S6).

#### Overlapping signatures of selection and ontology of genes bearing outlier

Overlapping signatures of selection from population divergence and GEA analyses were detected at genes 154_1 and QB_c10512 (Table 1, Fig. 5). The GEA analyses shared signatures of local adaptation in 12 additional genes (13 common SNPs). The population divergence analyses also shared signatures of local adaptation at one additional gene. Finally, 29 and six outlier SNPs were detected by *lfmm* and *Samβada* alone, respectively.

**Table 1.**
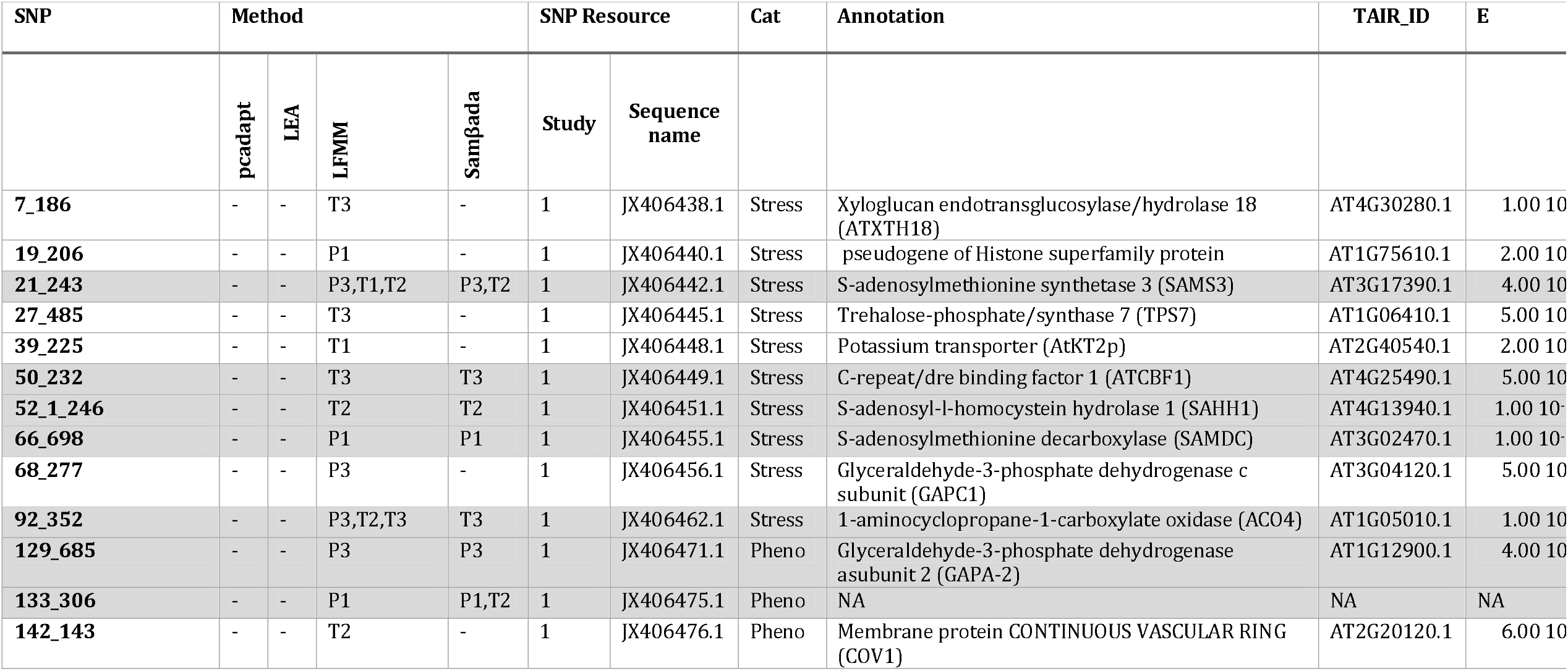
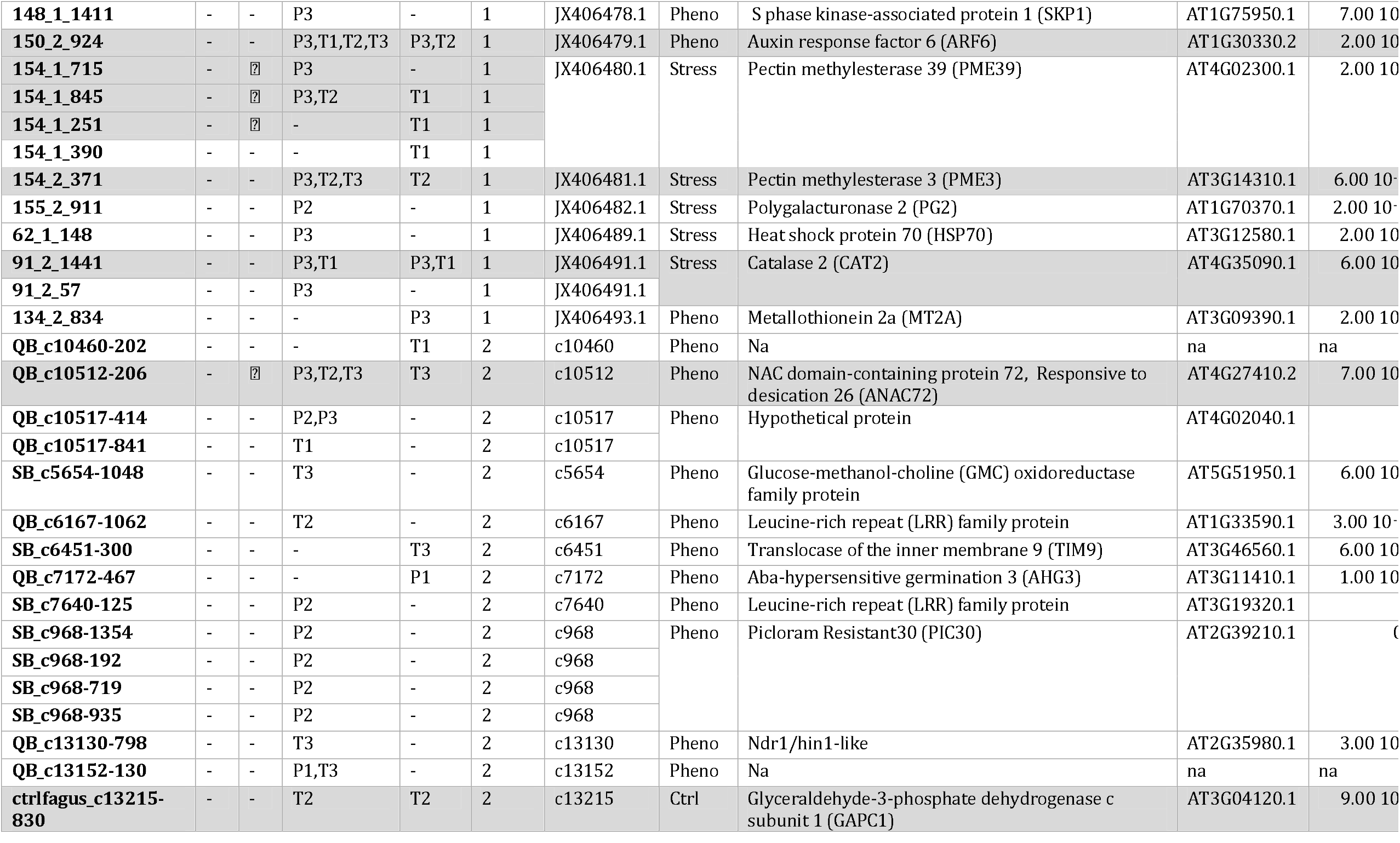
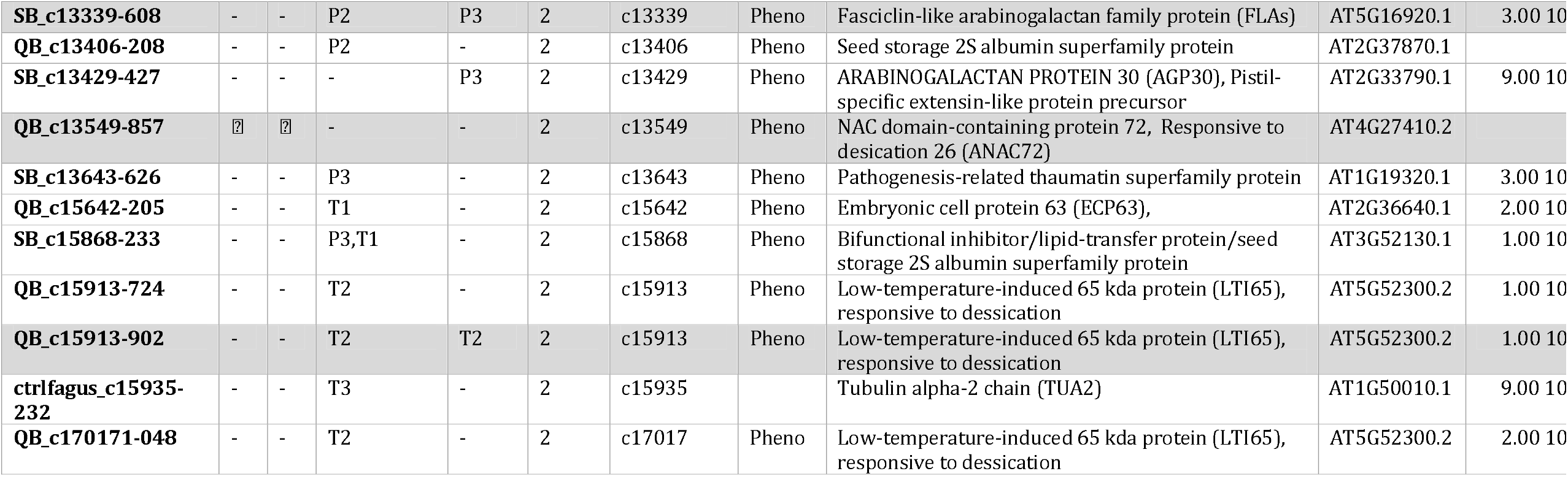
Outlier SNPs showing signature of divergent selection with at least one of the four methods (*pcadapt, lea, lfmm* and *Samβada*). We applied analysis-specific FDR cut-offs granting no expected false positives. For *lfmm* and *Samβada*, we report the climatic variable for which the genetic-environment association was found (T1-3: Temp1-3; P1-3: Precip 1-3). Converging selection signatures from at least two methods are underlined in grey. For each SNP, we code the study where it was described first (1: Lalagüe, et al. 2014 ; 2: Lesur, et al. 2015) and give the gene sequence (in Genbank for 1; in Lesur, et al. 2015 for 2). We finally provide for each candidate gene its category (stress-related, phenology-related or control genes), its annotation based on the homology with *Arabidopsis thaliana* sequence (with the TAIR ID and probability of matching E)

**Figure 5:**
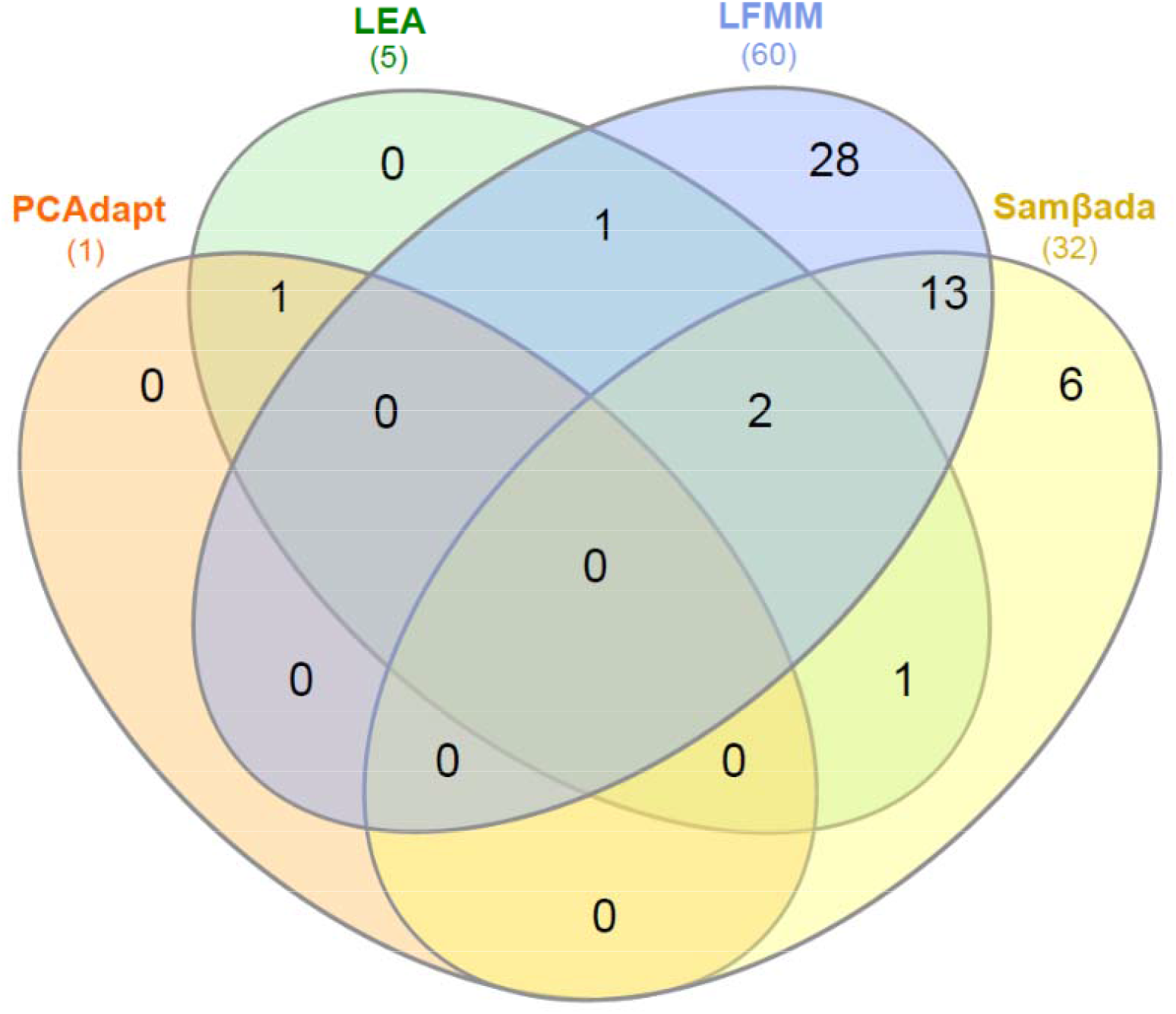
**Venn diagram of the private and common outliers identified by the different methods to detect signatures of divergent selection.**

While our panel of 270 SNPs included 67.9% of putatively phenology-related, 26.4% of stress-related and 5.7% of control-related genes, respectively, outliers included 61.4% of phenology-related, 34.1% of stress-related, and 4.5% of control genes (Table 1). Hence, there was no difference of category (stress, phenology, control) among initial and outlier genes (χ^2^ = 0.99, *p* = 0.61).

### Isolation by distance and environment

The variance partitioning and partial RDA analyses revealed a greater effect of spatial structure than of climatic structure on the spatial distribution of genetic variation (Figure S5, Table S7). Considering the 218 putatively neutral SNPs only, the climatic structure alone explained ∼1% of variance in the genetic structure, which is not statistically different from the null expectation of 0% variance explained (*F*_6,52_ = 1.17, *p* = 0.16). On the contrary, the contribution of the spatial structure alone was much larger (*R*^2^ = 16%) and its effect was statistically significant (*F*_5,52_ = 4.12, *p* < 0.001). The joint effect of climatic and spatial structures contributed significantly in explaining the genetic structure (*R*^2^ = 25%). In this case, such a joint effect is not equivalent to a standard interaction term, and relates to the intrinsic covariation of climatic and spatial effects.

When considering the set of 52 SNPs putatively under selection, the contribution of the climatic structure to the genetic structure was significant (*R*^2^ = 3%; *F*_6,52_ = 1.65, *p* = 0.006), but still smaller than that of the spatial structure (14%; *F*_5,52_ = 4.008, *p* < 0.001). The joint effects of climatic and spatial structures explained 32% of the genetic structure. Thus, the variance contributed by climatic and spatio-climatic structures combined was higher for the 52 outliers (∼35%) than for the supposedly neutral loci (∼26%; Fig. S5). Moreover, the temperature and precipitation components of the sole climatic effects explained a similar and statistically significant amount of variance (1.5% and 1.2%, respectively; *p* < 0.001; Table S7B) in the supposedly adaptive genetic structure. The variance contributed by shared spatial and temperature structures (21%) was higher than that contributed by shared spatial and precipitation structures (6%).

## Discussion

Our study supported our first expectation: we observed weak founder effects in beech, which indicates that past population demography is not likely to blur the detection of selection signatures. On the contrary, our second expectation was not met as we identified loci with the signature of divergent selection as often in genes involved in phenology as in genes involved in stress response. And counter to our third expectation, temperature and precipitation related variables were equally represented in the significant genotype-climate associations. Overall, our results suggest a balanced contribution of traits related with phenology and with stress responses to local adaptation in beech.

### Impact of past recolonization history on genetic diversity

Our results revealed a clear spatial disjunction between three main gene pools. A first pool (blue cluster) corresponds to the area harbouring beech glacial refugia in Southeastern Europe. Its geographical distribution matches well with results of genetic and palaeoecological studies showing that lineages from these refugia expanded only as far as the Northern Apennines and Central Carpathians (Leonardi & Menozzi, 1995; Magri, 2008; Magri et al., 2006). A second gene pool (red cluster) includes the Southwest European glacial refugia located on the Iberian Peninsula and in southern France, where beech persisted throughout several glacial cycles (de Lafontaine et al., 2014) and was even more abundant than in the southeastern refugia during the middle and upper Pleistocene (Magri et al., 2006). A third gene pool (green cluster) mostly corresponds to recently recolonized areas in Northern Europe (Sjölund, González-Díaz, Moreno-Villena, & Jump, 2017) and likely originates from glacial refugia located in the Eastern Alps-Slovenia and in Slovakia-Moravia (Magri, 2008). Boundaries between the three clusters were associated with strong admixture, which would be consistent with the relatively high number of recolonization routes known for beech as compared to other tree species (de Lafontaine et al. 2014; Magri, 2008).

Consistently with previous studies based on allozymes or microsatellites (Comps, Gomory, Letouzey, Thiebaut, & Petit, 2001; de Lafontaine et al., 2013), spatial patterns of SNP genetic diversity did not reflect signals of founder effects resulting from the post-glacial expansion. In particular, the northern populations (green cluster) showed values of Nei’s heterozygosity (*H*_*e*_) and genetic differentiation (*β*_*WT*_) similar to those of the southwestern populations (red cluster), while the southeastern populations (blue cluster) showed lower *H*_*e*_ and higher *β*_*WT*_. Hence, diversity across the 270 studied SNPs was higher, and differentiation lower, both in recently recolonized areas, and in areas where beech was more abundant in the past. These patterns likely result from the combination of several processes and life-history traits specific to trees in general and beech in particular: first, the long juvenile phase of forest trees strongly attenuates founder effects during colonization in a diffusive dispersal model (Austerlitz et al., 2000). Moreover, long-distance pollen dispersal is frequent in beech (Gauzere, Klein, & Oddou-Muratorio, 2013; Piotti et al., 2012), which is expected to increase the number of founders and the mixing of genes from distant sources, resulting in a rapid increase of genetic diversity after the initial colonization (Fayard et al., 2009; Lander, Klein, Roig, & Oddou-Muratorio, 2021; Paulose & Hallatschek, 2020). Finally, beech is one of the tree species that recolonised northern Europe the latest, and the factors that limited its ability to migrate probably also contributed to its retaining a high level of diversity along the expansion front (Roques et al., 2012; Saltré et al., 2013)

Range-wide spatial genetic structure (SGS) was statistically significant but weak. The strongest signal was found in the southeastern genetic cluster. This is consistent with the theoretical work of Slatkin (1993) on IBD, who showed that a species having restricted dispersal should exhibit SGS if enough time has elapsed after establishment. Since the south-eastern European populations (blue cluster) have undergone a relatively early and short-distance post-glacial expansion (Magri et al., 2006), they would have had the longest time for the establishment of SGS.

Altogether, our results hence agree on the absence of a marked signature of genetic drift and allele surfing in beech due to recolonization. It appears therefore warranted to assume that our analysis of genetic signatures of local adaptation is little burdened with such sources of uncertainty, in line with previous studies on temperate forest trees that have explicitly tested for such effects (Eckert et al., 2010; Ruiz Daniels et al., 2018; Temunović et al., 2020).

### Genomic signatures of local adaptation along climatic gradients

Two genes showed convergent signatures of selection using GEA and differentiation outlier analyses. GEA detected more outliers than differentiation outlier analyses, with 12 genes showing convergent signatures of divergent selection using *lfmm* and *Samβada*. The 52 outliers identified with at least one of the methods displayed significant IBE patterns (while the putatively neutral markers did not), consistently with the fact that allele frequencies co-vary with climatic variables at the loci under selection.

According to GEA, 50 associations were attributable to the temperature variables and 42 to the precipitation variables. Among the temperature variables, associations with climate continentality (22) were found more often than associations with mean temperature (12) or climate xericity (16). This finding may indicate that the risk of late frosts could represent a major constraint for the evolution of phenology-related traits in beech (Gauzere et al., 2020; Kreyling et al., 2014). Among the precipitation variables, associations with the coupling between precipitation and seasonal temperatures (25) were found more often than associations with precipitation abundance (8) or variability (9). This result could be due to genetic differentiation between locations where high precipitation occurs during the vegetation period (coupling) versus those where a precipitation deficit occurs during the vegetation period (decoupling). This would highlight the major role of low precipitation in driving patterns of local adaptation, in agreement with the known sensitivity of beech to drought (Aranda et al., 2015; Cuevo-Alarcon et al., 2021), and with the major role of maximal potential evapotranspiration as a driver of genetic differentiation for growth and survival (Gárate-Escamilla et al. 2019). Partial RDA analyses also indicate significant effects of temperature and precipitation on the genetic structure at all the 52 outlier loci together, even though the portion of genetic variance contributed by “pure” temperature or precipitation effects was low in both cases (1%). Considering that phenology-related candidate genes were slightly over-represented in our set of 270 SNPs, and assuming that optimal values of phenological traits are likely to vary primarily with temperature and photoperiod rather than with precipitation (Metcalf & Mitchell-Olds, 2009), the balanced contribution of precipitation and temperature variables to the genetic-climate associations suggests that range-wide local adaptation in beech is driven by traits related to various climate components (see also Garate-Escamilla et al. 2019).

Although we cannot completely rule out some false positives, the large number of outliers detected using GEA approaches is methodologically consistent. First, our sampling design with 64 populations covering beech range proves to be appropriate to account for steep ecological gradients, control for population structure, and ultimately optimize statistical power in GEA (Selmoni, Vajana, Guillaume, Rochat, & Joost, 2020). Moreover, the major post-glacial expansion axes of beech align with steep ecological gradients: the South-to-North axis of expansion opposes hot to cold climates, while the axis from Central Europe to Great Britain opposes continental to oceanic climates (Magri et al., 2006). Although the populations sampled in this study may not fully capture these axes of climate variation, such a configuration is expected to minimize the number of false positive with GEA, as compared to the opposite case where ecological gradients are orthogonal to the expansion axis (Frichot et al. 2015).

### Functional role of the genes under selection

Among the 139 candidate genes investigated, eighteen showed signatures of divergent selection with at least two methods. Eight of them had also shown signatures of divergent selection in previous studies (Table 1 and Table S8). Among these “best candidates”, the outlier gene 92 encodes for ACO, an enzyme involved in the production of the plant hormone ethylene, which regulates many plant developmental processes and stress responses. This study found a putative signature of selection at the non-synonymous locus at position 352, (coding for histidine or glutamine; Table 1), where the frequency of the homozygous genotype TT decreased with drought stress (Fig. 6a). This locus was also associated with annual and growth season temperatures in the study of Cuervo-Alarcon et al. (2018). Two other loci within this gene (although non-coding or synonymous) were also detected as outliers by Pluess et al. (2016), where their frequencies correlated with drought indices. These two previous studies were conducted in Switzerland, along drought and precipitation gradients using GEA approaches. Hence, our combined results suggest that the ACO gene could be under divergent selection at various hierarchical scales across Europe.

**Figure 6:**
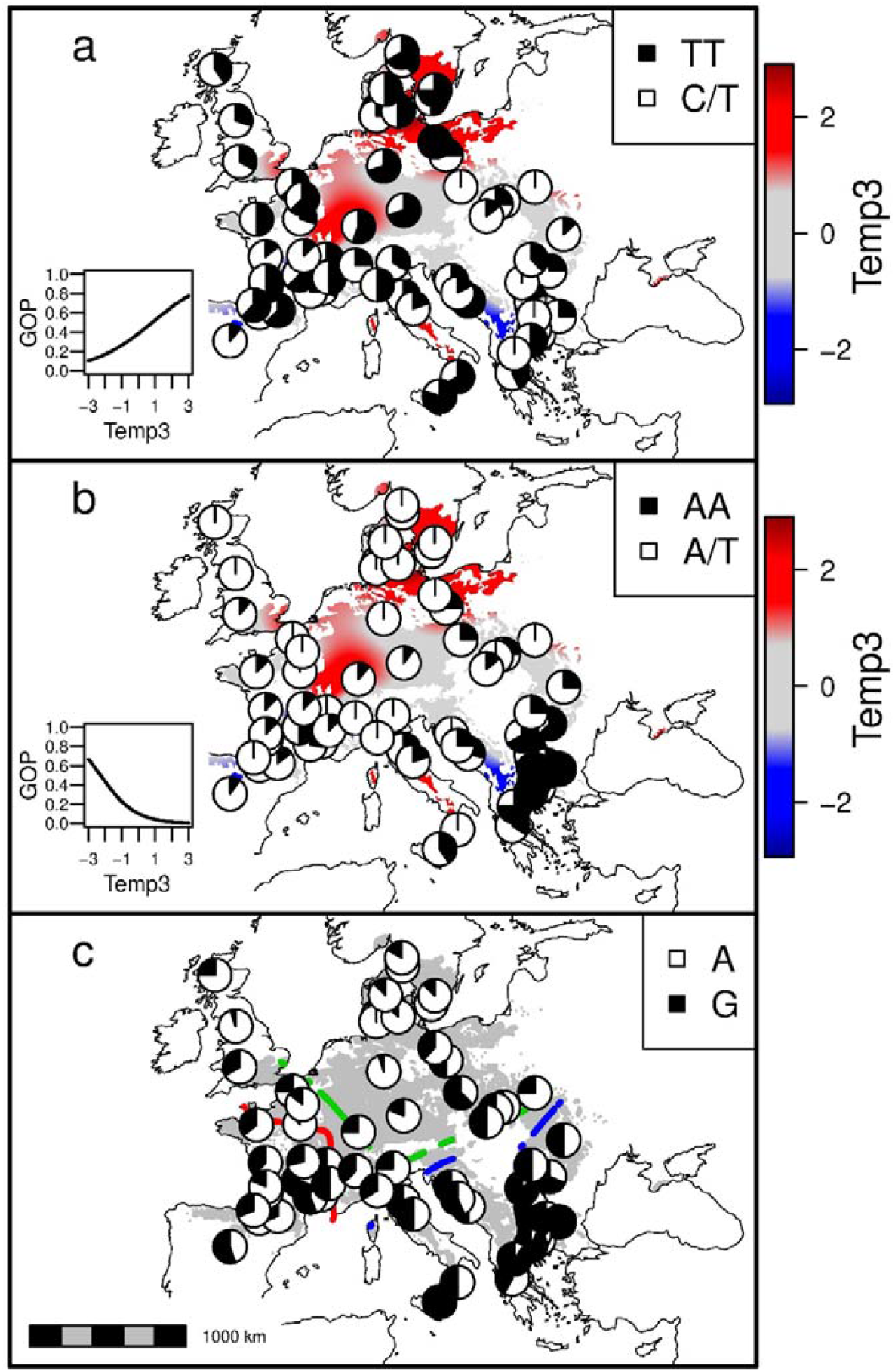
Variation in allelic/genotypic frequencies at three outlier SNPs (panel a: 92_352; b: QB_c10512; c: QB13549_857) across the studied geographic range. For panels (a) and (b), the graph on the left represents the predicted variation in genotype occurrence probability (GOP) across the environmental gradient as estimated by *Samβada*. Maps show the observed genotypic/allelic frequency superimposed on the environmental gradient (panels a and b) and spatial genetic structure (c).

Another interesting example is the outlier gene QB_c10512, which encodes for the NAC domain-containing protein 72, a transcription factor responsive to desiccation. This gene was found to be differentially expressed in beech quiescent buds by Lesur et al. (2015). At synonymous position 206 (in a Leucine-coding codon), the frequency of the AA genotype increased with drought stress (Fig. 6b). Two other variants within this gene (including a non-synonymous one) were also detected as outliers by Cuervo-Alarcon et al. (2018), where their frequencies also correlated with precipitation during the growing season. Moreover, the gene QB13549, detected as an outlier by *pcadapt* and *lea*, also encodes for the NAC domain-containing protein 72. At position 857, allele frequency showed a strong variation from Eastern to Western Europe (Fig. 6c).

The signatures of divergent selection were not particularly enriched in genes related to phenology. This is a counter-intuitive result, as previous quantitative genetic approaches failed to detect divergent selection at various physiological traits related to drought stress, but did detect significant differentiation in phenological (Gauzere et al., 2020; Hajek et al., 2016), growth and survival traits (Gárate-Escamilla et al., 2019; Gauzere et al., 2020). This is likely because the stress-related genes genotyped in this study are involved in the response to multiple stresses varying across climate gradients. Moreover, our panel of candidate genes probably determines a larger number of stress-related traits than usually phenotyped in quantitative genetic approaches (Gauzere et al., 2020; Hajek et al., 2016). Another limitation of quantitative genetic approaches is that they are usually conducted on a few populations and do hence not adequately cover the range-wide diversity of stress gradients (see also Garate-Escamilla et al. 2019). This comparison illustrates the complementarity of quantitative genetic and molecular approaches to investigate local adaptation (Rudman et al., 2018).

### Implications for conservation and management

The rates of expected (natural) species range shifts are likely to be insufficient for trees to track ongoing climate change (Saltré et al., 2013; Savolainen et al., 2007). In this context, there is an increasing interest in evolutionary-oriented management strategies, relying on the high genetic diversity observed within and among tree populations to adapt forest to ongoing climate change (Aitken & Whitlock, 2013; Lefèvre et al., 2014; Oney et al., 2013).

We showed that the spatial distribution of genetic diversity across beech range reflects both biogeographical history and adaptive processes. This has consequences for conservation, where the importance of maintaining adaptive genetic diversity - in addition to preserving as many lineages as possible - can not longer be overlooked (de Lafontaine et al. 2018; Ouborg, Pertoldi, Loeschcke, Bijlsma, & Hedrick, 2010; Shafer et al., 2015). While the conservation of lineages rely on the assessment of genetic boundaries, the conservation of adaptive diversity may require the identification of the relevant loci and the targeted conservation of specific alleles, genotypes or combinations thereof. Our results also have consequences for management, where knowledge of the genetic variants under selection, as combined with the estimation of their current spatial distribution and the prediction of future climate, may help inform decisions about assisted migration, perhaps under the form of an “enrichment” of existing stands with potentially favourable genotypes (Rellstab et al., 2016; Rochat, Selmoni, & Joost, 2021). This would however require a reliable validation of the adaptive meaning of our best candidates by independent proof, as well as the assessment of genotype⍰genotype interactions (to make sure there is no outbreeding depression) and of genotype⍰environment interactions (to avoid undesired, unforeseen under-performances of the introduced genotypes and their progeny in the new environments).

At the intersection of management and conservation lies the possibility to favour natural migration and regeneration dynamics, which could result in efficient mixing of genotypes in multiple environments, thus exposing them to natural selection and adaptive processes (Lefèvre et al., 2014). Here, the detailed analysis of barriers to gene flow is of the essence, to understand whether the barriers and “corridors” we detected have been caused by geographical features or isolation by adaptation: while in the former case it may be sensible to manage such barriers and corridors to shape gene flow, in the latter in may be difficult - or even detrimental - to force the modification of gene flow patterns.

## Acknowledgements

We thank Dalibor Ballian, Peter Zhelev and Aristotelis C. Papageorgiou for providing beech samples from Bosnia and Herzegovina, Bulgaria and Greece, as well as Ilaria Spanu, Mariaceleste Labriola, Catia Boggi and Adline Delcamp for technical assistance during lab work. SOM and AH were supported by the French National Research Agency Biodiversa project TipTree (ANR-12-EBID-0003); AP, FB and GGV supported by the Italian Ministry of University and Research (FOE-2019) under the projects “Climate Change” (CNR DTA.AD003.474) and “Green & Circular Economy” (CNR DTA.AD003.139); and AH by the French National Research Agency Cluster of Excellence project COTE (ANR-10-LABX-45, CLIMBEECH). The MassARRAY genotyping was performed at the Genome Transcriptome Facility of Bordeaux (grants from EU FP7 through Trees4Future research infrastructure, from the Conseil Régional d’Aquitaine n°20030304002FA and 20040305003FA, from the European Union FEDER n°2003227 and from Investissements d’Avenir ANR-10-EQPX-16-01).

## Data Accessibility

Raw SNPs genotypes, detailed information on SNPs and sampling locations, as well as R scripts for LD analyses, for differentiation outlier analyses (with *pcadapt*, and *lea*) and for GEA analyses (with *lfmm* and *Samβada*) will be made available at Portail Data INRAE: https://data.inrae.fr/, upon acceptance of the manuscript.

## Author Contributions

Conceptualization: S.O.M., G.G.V., D.P., E.V., A.P. Population sampling: D.P., F.B., G.G.V, A.H and F.P. Selection of candidate genes, SNPs and associated bioinformatics tools: D.P., I.L., G.L.P, S.O.M, G.G.V. SNP genotyping: D.P. and E.G. Statistical analyses: D.P., S.O.M, E.V., A.P. Writing-Original Draft : D.P., S.O.M, E.V., A.P. All the authors reviewed, edited and approved the final manuscript. Funding acquisition: S.O.M. and G.G.V.

