## Supplementary material for "Genetic signatures of divergent selection in European beech (*Fagus sylvatica L.*) are associated with the variation in temperature and precipitation across its distribution range": Suplementary figures and tables

### Table S1: Coordinates, country of origin and climatic characteristics of the 64 populations

We extracted the 19 bioclimatic variables from the WordClim 1.4 database (30 seconds resolution) over the period 1950-2000 and computed their average values. **MAT**: Mean Annual Temperature (°C) ; **TDR**: Mean diurnal range (°C); **isoT**: Isothermality (MAT/TAR ×100); **Tseas**: Temperature Seasonality (CV) ; **MaxTWarm**: Max Temperature of Warmest Period (°C); **MaxTCold**: Min Temperature of Coldest Period (°C); **TAR**: Temperature Annual Range (MaxTWarm-MaxTCold; °C); **MeanTWet**: Mean Temperature of Wettest Quarter (°C); **MeanTdry**: Mean Temperature of Driest Quarter (°C); **MeanTwarm**: Mean Temperature of Warmest Quarter (°C) ; **MeanTcold**: Mean Temperature of Coldest Quarter (°C); **MAP**: Annual Precipitation (mm); **Pwet**: Precipitation of Wettest Period (mm); **Pdry**: Precipitation of Driest Period (mm); **Pseas**: Precipitation Seasonality (CV); **PwetQ**: Precipitation of Wettest Quarter (mm); **PdryQ**: Precipitation of Driest Quarter (mm); **PwarmQ**: Precipitation of Warmest Quarter (mm); **PcoldQ**: Precipitation of Coldest Quarter (mm)

| Population | Long (°) | Lat (°) | Country | MAT | TDR | isoT | Tseas | MaxTWarm | MaxTCold | TAR | MeanTWet | MeanTdry | MeanTwarm | MeanTcold | MAP | Pwet | Pdry | Pseas | PwetQ | PdryQ | PwarmQ | PcoldQ |
|---|---|---|---|---|---|---|---|---|---|---|---|---|---|---|---|---|---|---|---|---|---|---|
| BG_58 | 25.5 | 42.6 | BG | 10.7 | 10.2 | 31.7 | 805.1 | 27.7 | -4.3 | 32 | 15.1 | 16.3 | 20.5 | 0.6 | 609 | 72 | 38 | 22.6 | 194 | 123 | 161 | 141 |
| BG_BN | 23.3 | 41.9 | BG | 3.9 | 9.1 | 33.4 | 665.0 | 18.6 | -8.5 | 27.1 | 6.5 | 9.3 | 12.0 | -4.1 | 693 | 73 | 33 | 21.9 | 205 | 120 | 165 | 190 |
| BG_VT | 23.3 | 42.6 | BG | 5.0 | 9.0 | 32.6 | 691.1 | 19.6 | -8.1 | 27.7 | 8.1 | 10.5 | 13.3 | -3.6 | 679 | 80 | 39 | 22.5 | 218 | 130 | 190 | 160 |
| BG-156 | 23.1 | 43.2 | BG | 8.2 | 9.5 | 31.8 | 751.6 | 23.7 | -6.1 | 29.8 | 15.3 | -0.1 | 17.1 | -1.4 | 669 | 84 | 42 | 25.6 | 229 | 133 | 200 | 140 |
| BG-157 | 23.1 | 43.2 | BG | 10.1 | 9.9 | 31.8 | 783.2 | 26.3 | -4.8 | 31.1 | 17.7 | 1.5 | 19.4 | 0.1 | 643 | 79 | 41 | 24.5 | 216 | 129 | 186 | 135 |
| BG-158 | 24.4 | 42.8 | BG | 6.8 | 9.3 | 31.5 | 753.0 | 22.3 | -7.4 | 29.7 | 14.1 | -1.8 | 15.9 | -2.6 | 700 | 93 | 41 | 28.0 | 245 | 139 | 216 | 146 |
| BG-16 | 23.7 | 41.5 | BG | 8.0 | 9.6 | 33.2 | 714.3 | 23.6 | -5.2 | 28.8 | 0.9 | 13.6 | 16.7 | -0.8 | 607 | 64 | 30 | 21.8 | 180 | 108 | 142 | 164 |
| BIH_Bih | 16.0 | 44.7 | BIH | 8.5 | 8.8 | 31.5 | 712.2 | 23.9 | -4.2 | 28.1 | 4.6 | 16.6 | 17.1 | -0.5 | 1187 | 140 | 78 | 20.8 | 392 | 248 | 259 | 305 |
| BIH_Cja | 17.5 | 43.5 | BIH | 4.9 | 7.0 | 28.8 | 648.7 | 18.6 | -5.6 | 24.2 | 1.8 | 12.6 | 13.0 | -2.8 | 1159 | 133 | 78 | 18.3 | 364 | 242 | 259 | 307 |
| BIH_Din | 16.6 | 44.0 | BIH | 6.2 | 7.4 | 28.8 | 677.3 | 20.4 | -5.1 | 25.5 | 2.8 | 14.1 | 14.6 | -2.1 | 1153 | 139 | 74 | 22.7 | 383 | 233 | 253 | 310 |

| Population | Long (°) | Lat (°) | Country | MAT | TDR | isoT | Tseas | MaxTWarm | MaxTCold | TAR | MeanTWet | MeanTdry | MeanTwarm | MeanTcold | MAP | Pwet | Pdry | Pseas | PwetQ | PdryQ | PwarmQ | PcoldQ |
|---|---|---|---|---|---|---|---|---|---|---|---|---|---|---|---|---|---|---|---|---|---|---|
| DE_07 | 7.8 | 48.0 | DE | 10.3 | 8.4 | 32.2 | 670.9 | 24.9 | -1.2 | 26.1 | 18.5 | 3.3 | 18.5 | 2.1 | 891 | 111 | 52 | 25.5 | 297 | 172 | 297 | 172 |
| DE_08 | 10.1 | 51.6 | DE | 7.8 | 8.3 | 33.7 | 638.0 | 21 | -3.7 | 24.7 | 15.5 | 0.6 | 15.5 | -0.2 | 769 | 87 | 49 | 19.1 | 245 | 158 | 245 | 177 |
| DE_09 | 11.8 | 48.9 | DE | 7.7 | 8.5 | 30.8 | 720.1 | 22.5 | -5 | 27.5 | 16.3 | 3.3 | 16.3 | -1.4 | 747 | 93 | 42 | 29.5 | 269 | 138 | 269 | 146 |
| DK_21 | 9.6 | 54.9 | DK | 8.2 | 6.0 | 27.6 | 604.6 | 19.7 | -2 | 21.7 | 9.3 | 3.2 | 15.6 | 1.0 | 774 | 86 | 41 | 22.9 | 241 | 136 | 211 | 175 |
| DK_24 | 10.2 | 55.3 | DK | 7.9 | 6.1 | 27.8 | 607.1 | 19.9 | -2.2 | 22.1 | 9.0 | 2.9 | 15.4 | 0.9 | 644 | 70 | 36 | 20.9 | 195 | 115 | 180 | 144 |
| DK_26 | 10.7 | 55.2 | DK | 8.0 | 5.1 | 24.1 | 611.0 | 19.3 | -1.7 | 21 | 9.4 | 2.6 | 15.5 | 0.9 | 605 | 66 | 34 | 21.2 | 180 | 107 | 173 | 118 |
| DK_32 | 9.4 | 54.5 | DK | 8.3 | 6.5 | 28.8 | 615.2 | 20.5 | -1.9 | 22.4 | 15.4 | 3.4 | 15.9 | 1.0 | 844 | 92 | 46 | 24.0 | 258 | 145 | 238 | 194 |
| DK_33 | 11.3 | 54.8 | DK | 8.2 | 5.1 | 23.1 | 629.0 | 19.3 | -2.8 | 22.1 | 15.7 | 2.6 | 15.9 | 0.8 | 606 | 65 | 35 | 18.8 | 181 | 116 | 176 | 122 |
| DK_37 | 10.2 | 56.1 | DK | 7.6 | 6.7 | 27.3 | 632.5 | 19.9 | -4.6 | 24.5 | 8.5 | 2.4 | 15.4 | 0.1 | 616 | 65 | 34 | 22.7 | 191 | 107 | 172 | 119 |
| ES_15 | -0.9 | 42.8 | ES | 8.9 | 9.8 | 39.5 | 558.1 | 22.7 | -2.2 | 24.9 | 3.2 | 15.9 | 15.9 | 2.3 | 994 | 107 | 55 | 16.6 | 290 | 199 | 199 | 272 |
| ES_16 | -3.5 | 41.1 | ES | 9.0 | 10.7 | 37.4 | 649.1 | 25.9 | -2.6 | 28.5 | 10.6 | 17.5 | 17.5 | 1.8 | 578 | 68 | 23 | 27.1 | 176 | 92 | 92 | 148 |
| FR_10 | 2.7 | 48.4 | FR | 10.7 | 8.7 | 36.1 | 595.9 | 24.2 | 0 | 24.2 | 16.4 | 7.0 | 18.0 | 3.3 | 668 | 64 | 46 | 8.5 | 178 | 147 | 170 | 166 |
| FR_12 | 5.0 | 46.0 | FR | 10.9 | 9.3 | 34.0 | 667.2 | 25.9 | -1.3 | 27.2 | 15.8 | 4.1 | 19.1 | 2.6 | 822 | 83 | 55 | 15.3 | 232 | 172 | 220 | 172 |
| FR_13 | 4.6 | 44.2 | FR | 12.5 | 10.0 | 35.8 | 646.7 | 28.1 | 0.2 | 27.9 | 13.1 | 20.6 | 20.6 | 4.7 | 792 | 103 | 34 | 25.8 | 263 | 147 | 147 | 189 |
| FR_AIG | 3.6 | 44.1 | FR | 7.0 | 8.8 | 36.2 | 563.3 | 20.8 | -3.6 | 24.4 | 11.7 | 1.0 | 14.1 | 0.5 | 913 | 91 | 57 | 14.3 | 250 | 203 | 219 | 215 |
| FR_ARG | 0.7 | 43.0 | FR | 10.3 | 9.7 | 38.5 | 585.1 | 24 | -1.2 | 25.2 | 12.3 | 4.4 | 17.5 | 3.2 | 919 | 95 | 61 | 14.2 | 265 | 202 | 234 | 207 |
| FR_AUB | 2.6 | 45.1 | FR | 5.9 | 9.2 | 38.4 | 552.7 | 19.3 | -4.6 | 23.9 | 7.6 | -0.1 | 12.7 | -0.6 | 1014 | 113 | 66 | 15.6 | 290 | 214 | 271 | 234 |
| FR_CHI | -0.4 | 46.1 | FR | 12.0 | 8.9 | 38.4 | 552.7 | 24.7 | 1.6 | 23.1 | 6.1 | 18.8 | 18.8 | 5.2 | 838 | 98 | 46 | 22.7 | 277 | 165 | 165 | 249 |
| FR_CIR | -0.3 | 44.4 | FR | 12.6 | 10.6 | 41.7 | 564.0 | 26.4 | 1.1 | 25.3 | 6.5 | 19.5 | 19.5 | 5.6 | 931 | 105 | 53 | 18.0 | 286 | 191 | 191 | 268 |
| FR_COL | 3.0 | 46.2 | FR | 9.4 | 9.9 | 38.8 | 590.5 | 23.4 | -2 | 25.4 | 14.9 | 5.5 | 16.7 | 2.2 | 772 | 93 | 47 | 23.0 | 238 | 147 | 228 | 160 |
| FR_FOU | -1.2 | 48.4 | FR | 10.6 | 8.1 | 39.0 | 495.0 | 22.2 | 1.4 | 20.8 | 5.5 | 16.6 | 16.6 | 4.6 | 768 | 86 | 49 | 18.0 | 239 | 163 | 163 | 217 |
| FR_HES | 2.0 | 50.4 | FR | 10.0 | 6.8 | 33.7 | 519.7 | 21 | 0.9 | 20.1 | 11.2 | 6.2 | 16.4 | 3.7 | 667 | 80 | 41 | 19.6 | 209 | 134 | 167 | 166 |
| FR_LAG | 2.9 | 49.6 | FR | 10.0 | 8.4 | 36.0 | 566.6 | 22.8 | -0.4 | 23.2 | 10.7 | 6.3 | 16.9 | 3.0 | 651 | 63 | 44 | 10.3 | 178 | 138 | 169 | 159 |

| Population | Long (°) | Lat (°) | Country | MAT | TDR | isoT | Tseas | MaxTWarm | MaxTCold | TAR | MeanTWet | MeanTdry | MeanTwarm | MeanTcold | MAP | Pwet | Pdry | Pseas | PwetQ | PdryQ | PwarmQ | PcoldQ |
|---|---|---|---|---|---|---|---|---|---|---|---|---|---|---|---|---|---|---|---|---|---|---|
| FR_LEO | 5.2 | 44.9 | FR | 6.0 | 8.5 | 33.4 | 617.3 | 20 | -5.4 | 25.4 | 2.7 | 13.4 | 13.7 | -1.5 | 1109 | 105 | 71 | 10.8 | 299 | 253 | 258 | 273 |
| FR_PAI | -0.3 | 44.7 | FR | 12.5 | 10.3 | 41.5 | 557.7 | 26.1 | 1.3 | 24.8 | 6.5 | 19.3 | 19.3 | 5.6 | 924 | 103 | 53 | 18.5 | 286 | 188 | 188 | 266 |
| FR_ROL | -1.4 | 43.3 | FR | 11.2 | 9.2 | 40.3 | 511.2 | 23.8 | 1 | 22.8 | 5.9 | 17.4 | 17.6 | 5.1 | 1305 | 157 | 67 | 21.8 | 421 | 249 | 255 | 395 |
| GB_01 | -4.8 | 57.3 | GB | 6.9 | 6.5 | 34.1 | 475.0 | 17.1 | -1.9 | 19 | 2.3 | 9.0 | 13.1 | 1.4 | 1402 | 169 | 75 | 29.5 | 478 | 228 | 267 | 433 |
| GB_02 | -3.0 | 54.3 | GB | 8.8 | 6.9 | 35.6 | 482.2 | 19.4 | -0.1 | 19.5 | 6.2 | 10.9 | 14.8 | 3.0 | 1014 | 109 | 58 | 23.7 | 324 | 189 | 234 | 267 |
| GB_03 | -2.7 | 51.8 | GB | 9.8 | 7.0 | 36.3 | 467.5 | 20.6 | 1.3 | 19.3 | 5.2 | 6.2 | 15.7 | 4.4 | 795 | 84 | 52 | 16.7 | 242 | 170 | 177 | 220 |
| GR_OX | 21.4 | 39.2 | GR | 12.5 | 11.0 | 36.2 | 694.9 | 29.6 | -0.7 | 30.3 | 5.7 | 21.2 | 21.2 | 4.2 | 930 | 141 | 21 | 51.2 | 379 | 87 | 87 | 362 |
| GR_PO | 23.0 | 41.2 | GR | 12.2 | 10.3 | 33.1 | 768.9 | 29 | -2 | 31 | 4.2 | 21.2 | 21.6 | 2.8 | 491 | 58 | 27 | 22.4 | 148 | 90 | 105 | 128 |
| GR_TP | 21.6 | 40.4 | GR | 6.6 | 10.3 | 34.8 | 690.7 | 23.1 | -6.6 | 29.7 | -0.4 | 14.8 | 15.1 | -1.8 | 788 | 94 | 39 | 27.2 | 259 | 126 | 133 | 229 |
| IT_16 | 7.6 | 45.6 | IT | -0.2 | 6.1 | 28.1 | 575.9 | 11.5 | -10.3 | 21.8 | 1.3 | 7.0 | 7.0 | -6.7 | 1726 | 160 | 130 | 7.1 | 459 | 411 | 411 | 431 |
| IT_18 | 11.8 | 43.8 | IT | 10.1 | 7.0 | 28.2 | 659.2 | 24.1 | -0.8 | 24.9 | 6.7 | 18.4 | 18.4 | 2.4 | 886 | 109 | 47 | 22.8 | 288 | 165 | 165 | 218 |
| IT_19 | 10.9 | 45.8 | IT | 5.6 | 7.3 | 28.4 | 663.7 | 19.6 | -6.1 | 25.7 | 13.6 | -2.4 | 13.8 | -2.4 | 752 | 94 | 30 | 37.6 | 261 | 96 | 258 | 96 |
| IT_19P | 9.5 | 44.5 | IT | 6.6 | 5.7 | 25.1 | 618.1 | 19.2 | -3.6 | 22.8 | 7.8 | -0.2 | 14.3 | -0.6 | 980 | 120 | 58 | 23.1 | 325 | 202 | 208 | 207 |
| IT_20 | 16.6 | 39.0 | IT | 14.1 | 6.8 | 30.2 | 583.7 | 26.8 | 4.4 | 22.4 | 11.9 | 21.4 | 21.5 | 7.6 | 932 | 143 | 18 | 58.5 | 394 | 67 | 95 | 363 |
| IT_32 | 12.6 | 43.1 | IT | 11.1 | 7.4 | 28.8 | 670.6 | 25.6 | 0 | 25.6 | 12.2 | 19.6 | 19.6 | 3.3 | 905 | 108 | 52 | 20.4 | 287 | 191 | 191 | 207 |
| IT_R | 15.0 | 37.8 | IT | 9.8 | 6.2 | 26.5 | 641.5 | 23.2 | -0.1 | 23.3 | 7.2 | 18.0 | 18.0 | 2.7 | 667 | 91 | 16 | 49.3 | 258 | 60 | 60 | 235 |
| PL_115 | 20.6 | 49.5 | PL | 4.6 | 8.7 | 30.0 | 743.2 | 19.3 | -9.7 | 29 | 13.5 | -4.0 | 13.5 | -4.8 | 934 | 137 | 43 | 42.1 | 377 | 143 | 377 | 150 |
| PL_117 | 16.9 | 50.3 | PL | 6.8 | 8.4 | 30.9 | 707.4 | 21.1 | -6.2 | 27.3 | 15.2 | -1.1 | 15.2 | -2.2 | 639 | 91 | 28 | 48.2 | 271 | 86 | 271 | 89 |

| Population | Long (°) | Lat (°) | Country | MAT | TDR | isoT | Tseas | MaxTWarm | MaxTCold | TAR | MeanTWet | MeanTdry | MeanTwarm | MeanTcold | MAP | Pwet | Pdry | Pseas | PwetQ | PdryQ | PwarmQ | PcoldQ |
|---|---|---|---|---|---|---|---|---|---|---|---|---|---|---|---|---|---|---|---|---|---|---|
| PL_121 | 15.6 | 52.2 | PL | 8.8 | 7.8 | 27.1 | 780.1 | 23.9 | -4.7 | 28.6 | 18.1 | 0.1 | 18.1 | -1.1 | 560 | 70 | 29 | 28.9 | 196 | 94 | 196 | 109 |
| PL_66 | 23.4 | 50.3 | PL | 7.1 | 8.2 | 27.5 | 799.8 | 22.4 | -7.4 | 29.8 | 16.5 | -2.0 | 16.5 | -3.1 | 627 | 87 | 29 | 40.6 | 245 | 93 | 245 | 98 |
| PL_68 | 14.6 | 53.0 | PL | 8.6 | 7.8 | 27.9 | 757.7 | 23.3 | -4.6 | 27.9 | 17.6 | 0.1 | 17.6 | -1.1 | 551 | 69 | 30 | 26.2 | 188 | 99 | 188 | 110 |
| PL_69 | 20.0 | 49.3 | PL | 5.2 | 9.1 | 30.8 | 749.5 | 20.2 | -9.3 | 29.5 | 14.0 | -3.4 | 14.0 | -4.4 | 1040 | 166 | 44 | 48.2 | 442 | 141 | 442 | 147 |
| RO_CA | 24.7 | 45.3 | RO | 6.4 | 9.8 | 32.0 | 769.4 | 21.6 | -8.9 | 30.5 | 14.0 | -2.2 | 15.5 | -3.5 | 764 | 115 | 39 | 42.8 | 311 | 120 | 299 | 123 |
| RO_DB | 22.3 | 44.6 | RO | 10.4 | 9.4 | 31.0 | 798.6 | 26.8 | -3.6 | 30.4 | 18.2 | 1.8 | 19.8 | 0.2 | 658 | 90 | 41 | 28.0 | 232 | 124 | 213 | 135 |
| RO_GH | 25.9 | 47.5 | RO | 6.5 | 9.4 | 30.2 | 820.1 | 22.1 | -9.1 | 31.2 | 14.8 | -4.0 | 16.2 | -4.0 | 657 | 107 | 28 | 54.2 | 293 | 87 | 283 | 87 |
| RO_SC | 23.1 | 46.0 | RO | 7.3 | 9.2 | 32.1 | 731.6 | 22.1 | -6.5 | 28.6 | 14.4 | -1.0 | 16.0 | -2.0 | 770 | 118 | 40 | 41.6 | 309 | 123 | 297 | 132 |
| SE_04 | 11.7 | 57.9 | SE | 7.3 | 6.2 | 25.9 | 686.8 | 19.8 | -4.3 | 24.1 | 12.7 | 1.6 | 15.7 | -1.0 | 695 | 81 | 36 | 27.2 | 231 | 115 | 194 | 149 |
| SE_05 | 14.3 | 55.6 | SE | 7.7 | 5.5 | 24.8 | 636.0 | 19.7 | -2.4 | 22.1 | 15.3 | 2.2 | 15.7 | 0.4 | 591 | 64 | 33 | 21.1 | 180 | 105 | 168 | 139 |
| SE_25 | 11.6 | 58.3 | SE | 6.9 | 6.0 | 24.9 | 687.8 | 19.6 | -4.5 | 24.1 | 12.1 | 1.2 | 15.5 | -1.3 | 746 | 86 | 39 | 26.5 | 244 | 125 | 207 | 162 |
| SE_28 | 14.6 | 56.1 | SE | 7.3 | 6.0 | 26.0 | 650.6 | 20.1 | -3 | 23.1 | 15.0 | 1.9 | 15.6 | -0.2 | 610 | 69 | 35 | 21.6 | 188 | 110 | 174 | 144 |
| SK_23 | 19.1 | 48.6 | SK | 8.7 | 10.7 | 32.7 | 803.5 | 25.4 | -7.2 | 32.6 | 18.0 | 0.0 | 18.0 | -1.8 | 681 | 85 | 41 | 24.2 | 220 | 124 | 220 | 137 |

*Country : BG= Bulgaria; BIH= Bosnia and Herzegovina; DE= Germany; DK= Denmark; ES= Spain; FR=France; GB= Great Britain; IT=Italy; PL=Poland; RO= Romania; SE= Sweden; SK= Slovakia.

### Table S2: Diversity parameters and cluster assignments of the 64 populations.

**q1, q2, q2:** q-values for the assignation to cluster C1, C2, C3 (respectively green, red and blue on Fig. 2) using DAPC; **n:** number of genotyped individuals; **Na**: mean number of alleles; **Ar**: allelic richness; **%polloc:** percentage of polymorphic loci**; He**: expected heterozygosity; **Ho**: observed heterozygosity; **$F_{IS}$**: inbreeding coefficient. **βwt** : genetic differentiation from the entire pool.

| Population | q1 | q2 | q3 | n | Na | Ar | %polloc | $H_E$ | $H_0$ | $F_{IS}$ | $β_{WT}$ |
|---|---|---|---|---|---|---|---|---|---|---|---|
| BG_156 | 0.15 | 0.19 | 0.66 | 4 | 1.67 | 1.51 | 67 | 0.287 | 0.277 | 0.005 | 0.102 |
| BG_157 | 0.42 | 0.10 | 0.47 | 4 | 1.71 | 1.56 | 71 | 0.306 | 0.297 | -0.001 | 0.039 |
| BG_158 | 0.12 | 0.18 | 0.70 | 3 | 1.67 | 1.56 | 67 | 0.316 | 0.333 | -0.100 | -0.007 |
| BG_16 | 0.05 | 0.06 | 0.88 | 4 | 1.65 | 1.51 | 65 | 0.265 | 0.272 | -0.045 | 0.162 |
| BG_58 | 0.06 | 0.12 | 0.83 | 4 | 1.71 | 1.57 | 71 | 0.310 | 0.298 | 0.003 | 0.032 |
| BG_BN | 0.12 | 0.12 | 0.76 | 8 | 1.83 | 1.59 | 83 | 0.298 | 0.313 | -0.045 | 0.057 |
| BG_VT | 0.05 | 0.06 | 0.89 | 7 | 1.77 | 1.56 | 77 | 0.288 | 0.291 | -0.011 | 0.093 |
| BIH_Bih | 0.29 | 0.07 | 0.64 | 8 | 1.75 | 1.56 | 75 | 0.291 | 0.327 | -0.116 | 0.078 |
| BIH_Cja | 0.11 | 0.06 | 0.83 | 7 | 1.75 | 1.56 | 75 | 0.295 | 0.304 | -0.047 | 0.070 |
| BIH_Din | 0.14 | 0.09 | 0.77 | 8 | 1.77 | 1.55 | 77 | 0.280 | 0.291 | -0.026 | 0.115 |
| DE_07 | 0.52 | 0.37 | 0.11 | 9 | 1.81 | 1.6 | 81 | 0.306 | 0.307 | -0.008 | 0.037 |
| DE_08 | 0.74 | 0.10 | 0.16 | 10 | 1.81 | 1.58 | 81 | 0.302 | 0.309 | -0.007 | 0.047 |
| DE_09 | 0.65 | 0.11 | 0.24 | 10 | 1.83 | 1.59 | 83 | 0.300 | 0.289 | 0.030 | 0.058 |
| DK_21 | 0.80 | 0.15 | 0.05 | 4 | 1.72 | 1.58 | 72 | 0.319 | 0.318 | -0.022 | -0.006 |
| DK_24 | 0.79 | 0.12 | 0.09 | 4 | 1.73 | 1.58 | 73 | 0.318 | 0.347 | -0.106 | -0.019 |
| DK_26 | 0.85 | 0.08 | 0.07 | 4 | 1.67 | 1.52 | 67 | 0.302 | 0.262 | 0.092 | 0.065 |
| DK_32 | 0.73 | 0.16 | 0.11 | 4 | 1.69 | 1.53 | 69 | 0.294 | 0.286 | -0.004 | 0.076 |
| DK_33 | 0.86 | 0.06 | 0.09 | 4 | 1.68 | 1.56 | 68 | 0.298 | 0.316 | -0.080 | 0.053 |
| DK_37 | 0.82 | 0.08 | 0.10 | 4 | 1.71 | 1.59 | 71 | 0.319 | 0.339 | -0.083 | -0.012 |
| ES_15 | 0.10 | 0.84 | 0.06 | 10 | 1.84 | 1.6 | 84 | 0.297 | 0.299 | -0.001 | 0.065 |
| ES_16 | 0.07 | 0.78 | 0.15 | 10 | 1.85 | 1.6 | 85 | 0.297 | 0.289 | 0.022 | 0.065 |
| FR_10 | 0.40 | 0.43 | 0.17 | 10 | 1.85 | 1.6 | 85 | 0.301 | 0.312 | -0.027 | 0.050 |
| FR_12 | 0.26 | 0.66 | 0.08 | 9 | 1.83 | 1.62 | 83 | 0.316 | 0.305 | 0.029 | 0.005 |
| FR_13 | 0.52 | 0.37 | 0.12 | 8 | 1.82 | 1.6 | 82 | 0.304 | 0.298 | 0.026 | 0.045 |
| FR_AIG | 0.09 | 0.63 | 0.28 | 8 | 1.82 | 1.61 | 82 | 0.307 | 0.304 | 0.002 | 0.034 |
| FR_ARG | 0.07 | 0.84 | 0.09 | 8 | 1.86 | 1.62 | 86 | 0.307 | 0.316 | -0.028 | 0.031 |
| FR_AUB | 0.11 | 0.76 | 0.13 | 8 | 1.86 | 1.62 | 86 | 0.313 | 0.313 | 0.002 | 0.015 |
| FR_CHI | 0.07 | 0.68 | 0.25 | 8 | 1.84 | 1.62 | 84 | 0.306 | 0.326 | -0.052 | 0.033 |
| FR_CIR_B | 0.14 | 0.74 | 0.13 | 8 | 1.86 | 1.61 | 86 | 0.322 | 0.311 | 0.016 | -0.011 |
| FR_COL | 0.18 | 0.71 | 0.12 | 8 | 1.87 | 1.65 | 87 | 0.322 | 0.337 | -0.041 | -0.018 |
| FR_FOU | 0.33 | 0.44 | 0.22 | 8 | 1.85 | 1.62 | 85 | 0.314 | 0.328 | -0.029 | 0.006 |
| FR_HES | 0.39 | 0.45 | 0.17 | 8 | 1.83 | 1.65 | 83 | 0.327 | 0.336 | -0.027 | -0.034 |
| FR_LAG | 0.38 | 0.44 | 0.18 | 8 | 1.81 | 1.61 | 81 | 0.301 | 0.313 | -0.035 | 0.049 |
| FR_LEO | 0.14 | 0.74 | 0.12 | 8 | 1.85 | 1.64 | 85 | 0.319 | 0.318 | -0.003 | -0.006 |
| FR_PAI | 0.11 | 0.80 | 0.09 | 8 | 1.86 | 1.62 | 86 | 0.312 | 0.307 | 0.021 | 0.017 |
| FR_ROL | 0.05 | 0.91 | 0.04 | 8 | 1.80 | 1.55 | 80 | 0.290 | 0.283 | 0.016 | 0.087 |

| Population | q1 | q2 | q3 | n | Na | Ar | %polloc | $H_E$ | $H_0$ | $F_{IS}$ | $\beta_{WT}$ |
|---|---|---|---|---|---|---|---|---|---|---|---|
| GB_01 | 0.35 | 0.49 | 0.16 | 10 | 1.85 | 1.61 | 85 | 0.307 | 0.282 | 0.062 | 0.038 |
| GB_02 | 0.50 | 0.44 | 0.06 | 9 | 1.83 | 1.59 | 83 | 0.295 | 0.307 | -0.028 | 0.068 |
| GB_03 | 0.49 | 0.42 | 0.09 | 9 | 1.85 | 1.6 | 85 | 0.301 | 0.306 | -0.011 | 0.050 |
| GR_OX | 0.09 | 0.08 | 0.83 | 7 | 1.80 | 1.59 | 80 | 0.306 | 0.296 | 0.013 | 0.042 |
| GR_PO | 0.06 | 0.06 | 0.88 | 7 | 1.80 | 1.57 | 81 | 0.296 | 0.282 | 0.035 | 0.073 |
| GR_TP | 0.05 | 0.06 | 0.90 | 5 | 1.73 | 1.54 | 73 | 0.291 | 0.302 | -0.042 | 0.081 |
| IT_16 | 0.36 | 0.28 | 0.36 | 4 | 1.76 | 1.59 | 76 | 0.317 | 0.321 | -0.031 | -0.001 |
| IT_18 | 0.29 | 0.07 | 0.64 | 6 | 1.73 | 1.54 | 73 | 0.285 | 0.301 | -0.061 | 0.098 |
| IT_19 | 0.63 | 0.12 | 0.25 | 6 | 1.77 | 1.58 | 77 | 0.292 | 0.315 | -0.070 | 0.074 |
| IT_20 | 0.04 | 0.07 | 0.90 | 9 | 1.80 | 1.56 | 80 | 0.280 | 0.290 | -0.033 | 0.116 |
| IT_32 | 0.25 | 0.15 | 0.61 | 5 | 1.73 | 1.57 | 73 | 0.307 | 0.312 | -0.032 | 0.030 |
| IT_P | 0.46 | 0.10 | 0.45 | 4 | 1.69 | 1.54 | 69 | 0.293 | 0.281 | 0.005 | 0.082 |
| IT_R | 0.03 | 0.06 | 0.91 | 5 | 1.63 | 1.49 | 63 | 0.251 | 0.289 | -0.136 | 0.195 |
| PL_115 | 0.49 | 0.23 | 0.28 | 4 | 1.62 | 1.48 | 62 | 0.248 | 0.293 | -0.167 | 0.201 |
| PL_117 | 0.50 | 0.07 | 0.43 | 4 | 1.71 | 1.57 | 71 | 0.296 | 0.308 | -0.056 | 0.062 |
| PL_121 | 0.78 | 0.07 | 0.15 | 4 | 1.71 | 1.57 | 71 | 0.301 | 0.327 | -0.097 | 0.041 |
| PL_66 | 0.61 | 0.07 | 0.32 | 4 | 1.70 | 1.55 | 70 | 0.302 | 0.293 | -0.003 | 0.054 |
| PL_68 | 0.59 | 0.14 | 0.27 | 4 | 1.75 | 1.59 | 75 | 0.310 | 0.317 | -0.045 | 0.021 |
| PL_69 | 0.61 | 0.08 | 0.31 | 4 | 1.71 | 1.59 | 71 | 0.317 | 0.330 | -0.060 | -0.006 |
| RO_CA | 0.22 | 0.09 | 0.69 | 8 | 1.79 | 1.59 | 79 | 0.307 | 0.320 | -0.034 | 0.057 |
| RO_DB | 0.29 | 0.06 | 0.65 | 8 | 1.81 | 1.61 | 81 | 0.305 | 0.309 | -0.014 | 0.029 |
| RO_GH | 0.14 | 0.10 | 0.76 | 7 | 1.82 | 1.59 | 82 | 0.287 | 0.272 | 0.047 | 0.039 |
| RO_SC | 0.41 | 0.06 | 0.53 | 8 | 1.78 | 1.56 | 78 | 0.300 | 0.294 | 0.010 | 0.099 |
| SE_04 | 0.87 | 0.09 | 0.05 | 10 | 1.76 | 1.57 | 76 | 0.285 | 0.285 | 0.000 | 0.103 |
| SE_05 | 0.90 | 0.05 | 0.06 | 10 | 1.83 | 1.61 | 83 | 0.300 | 0.308 | -0.025 | 0.055 |
| SE_25 | 0.84 | 0.10 | 0.06 | 3 | 1.63 | 1.52 | 63 | 0.287 | 0.297 | -0.071 | 0.090 |
| SE_28 | 0.83 | 0.06 | 0.11 | 4 | 1.69 | 1.55 | 69 | 0.294 | 0.290 | -0.011 | 0.074 |
| SK_23 | 0.57 | 0.10 | 0.33 | 7 | 1.77 | 1.58 | 77 | 0.304 | 0.319 | -0.042 | 0.037 |

### Figure S1: Patterns of population structure and admixture inferred by Discriminant Analysis of Principal Components (DAPC)

(a) Variation in the Bayesian Information Criterion (BIC) with the possible number of clusters
(b) PCA on allelic frequencies
(c) Geographical distribution of the frequency of each cluster in each population. Each point correspond to one individual. We used a simple jitter around population coordinates to avoid superposition.

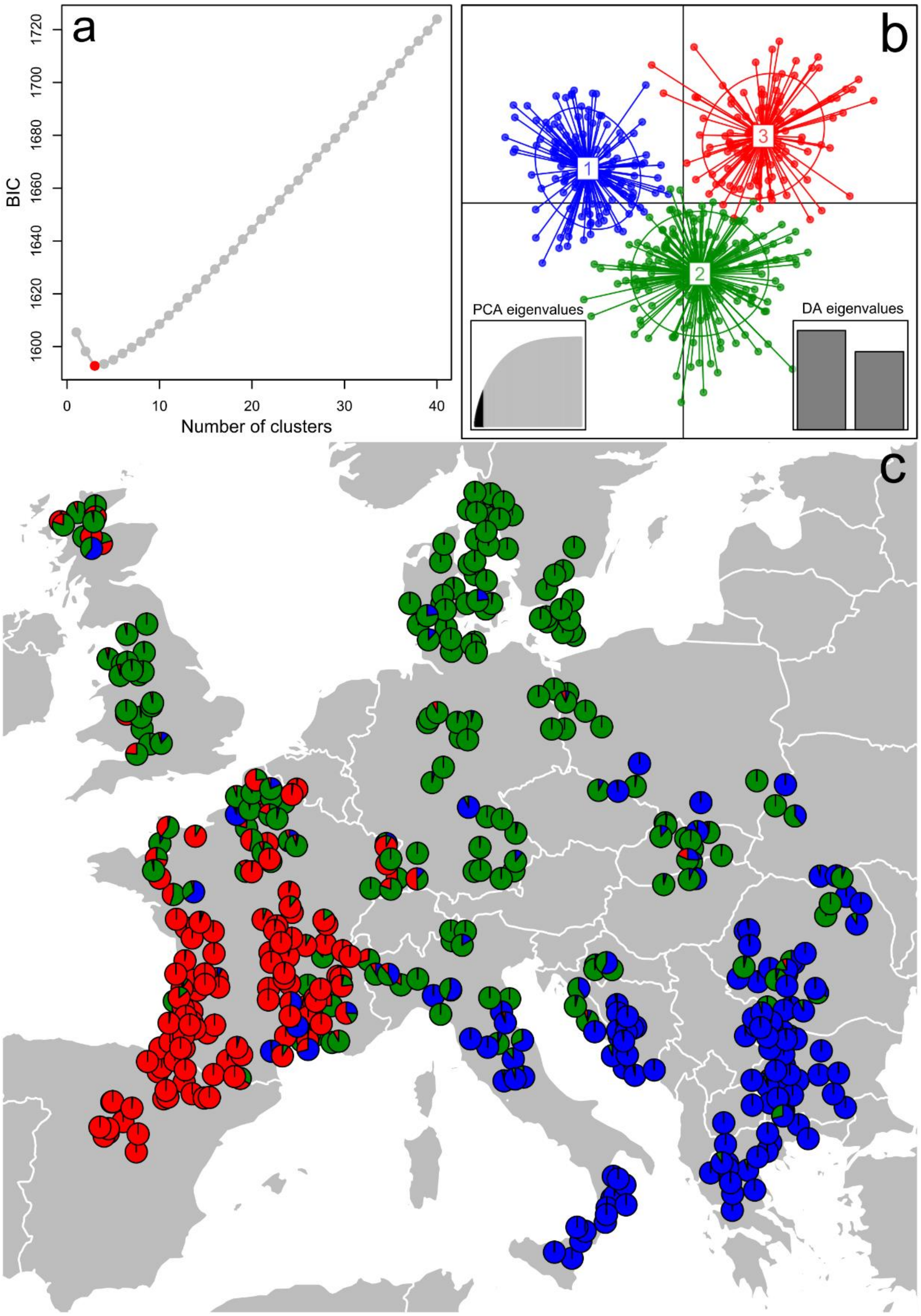

### Figure S2: Variation of genetic diversity and differentiation among populations assigned to different genetic clusters.

Boxplots were used to visualize the variation in (a) expected heterozygosity, He, (b) percentage of polymorphic loci,%polloc and (c) genetic differentiation relative to the entire pool,$\beta_{WT}$, between the three inferred genetic clusters. These analyses were run on the 47 populations assigned to the green (16), red (12) and blue clusters as defined by STRUCTURE and DAPC. We used Tukey test to compare means of each variable among each pairs of clusters.

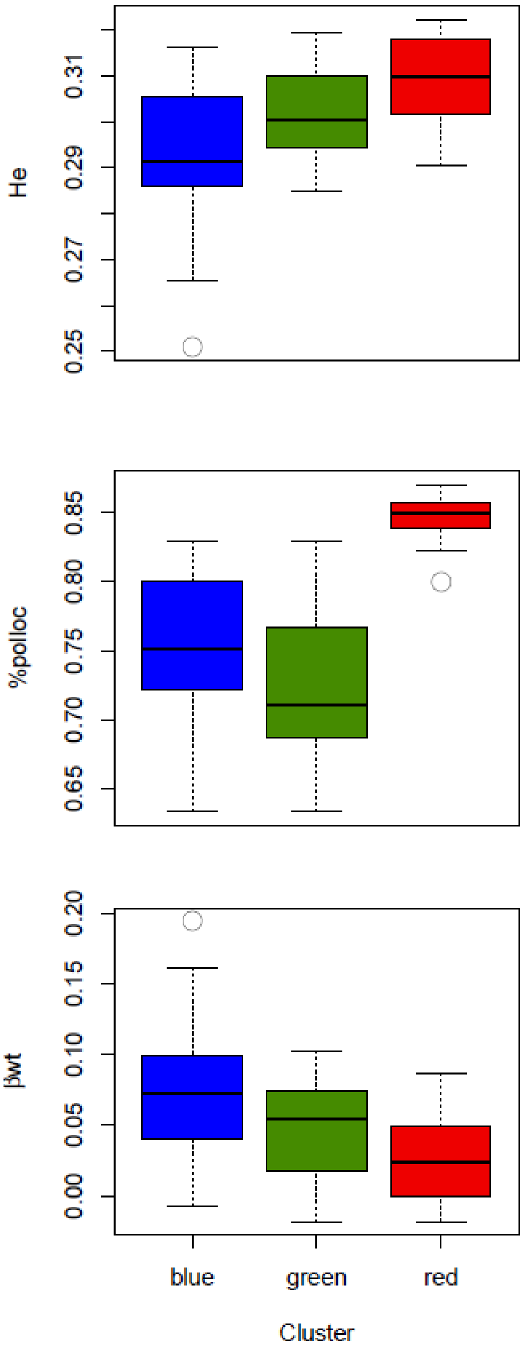

### Figure S3: Patterns of variation in the percentage of polymorphic SNPs within a population for the two different SNP arrays used in this study.

The first array of SNP markers ("KASP array"), developed by Lalagüe et al. (2014), was based on the genotyping of one population south-eastern France, while the second array of SNP markers ("Sequenom Array"), developed for this study, was based on a larger number of populations representative of beech distribution range. The fraction of polymorphic loci displayed below for each SNP array and their combination suggests that that diversity patterns do not markedly differ between the two arrays.

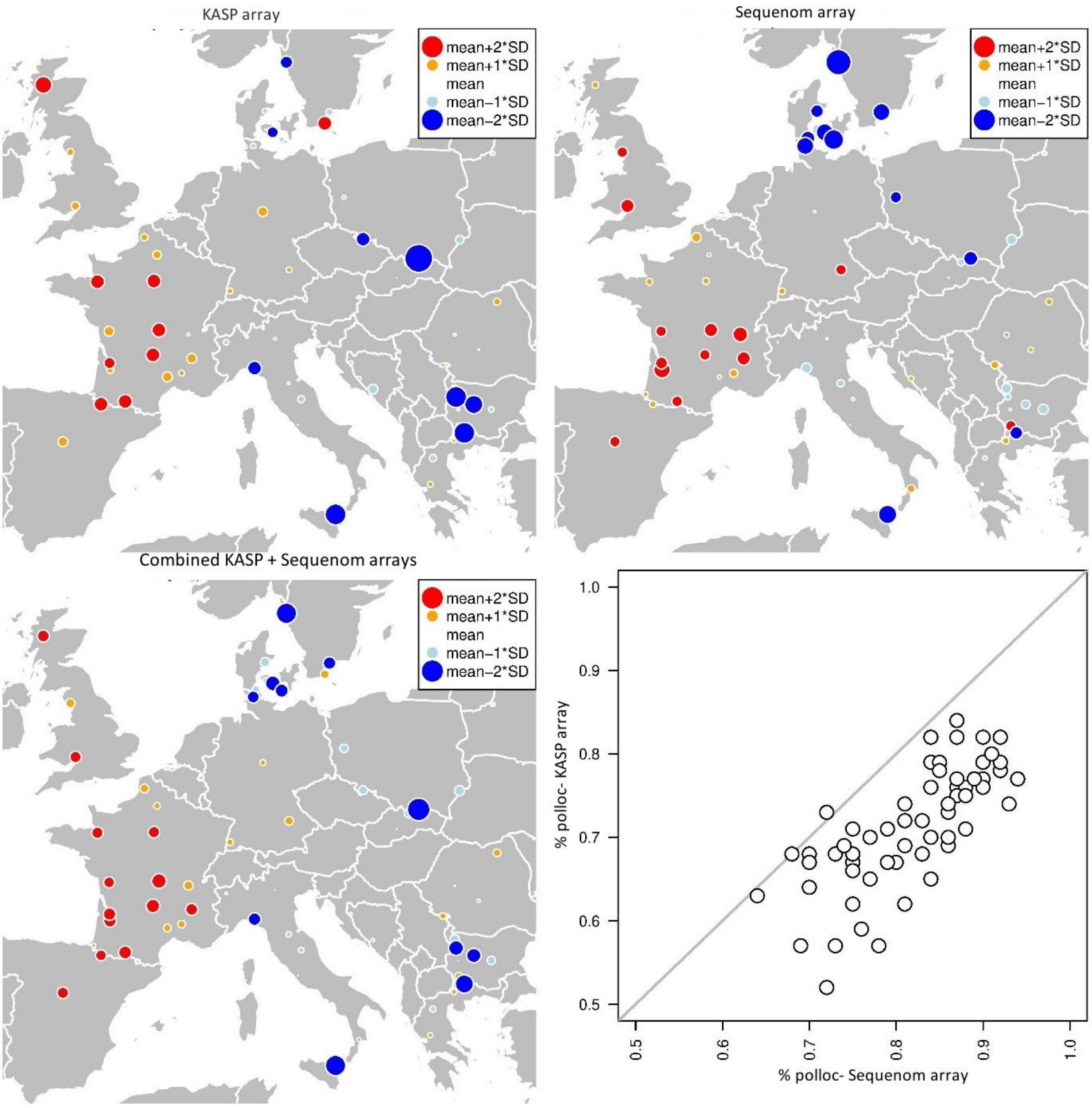

### Table S3: Signal of isolation by distance (IBD) on genetic differentiation between all populations, and between populations belonging to each cluster.

**nPop/nIndiv:** number of populations/individuals considered in the SGS analyses; **blog**: slope of the regression of $F_{ST}/1\text{-}F_{ST}$ values against logarithm of distance with its standard error (**se**); **bmin, bmax:** distribution envelope of blog values under the hypothesis of complete spatial randomness; **p-value** associated to the one-sided test with H1 $blog_{observed} > blog_{expected}$.

| *Group* | *nPop* | *nIndiv* | *blog* | *se* | *bmin* | *bmax* | *pvalue* |
|---|---|---|---|---|---|---|---|
| *Blue cluster* | 29 | 117 | 0.0192 | 0.0043 | -0.0140 | 0.0153 | 0.0056 |
| *Red cluster* | 20 | 99 | 0.0030 | 0.0057 | -0.0093 | 0.0095 | 0.2685 |
| *Green cluster* | 36 | 142 | 0.0079 | 0.0024 | -0.0086 | 0.0094 | 0.05 |
| *All populations* | 64 | 430 | 0.0266 | 0.0021 | -0.0030 | 0.0033 | 0 |

### Figure S4: Patterns of spatial genetic structure among all the 64 studied populations (black) and within each DAPC-defined cluster (in red, blue and green).

The values of genetic differentiation (measured by $F_{ST}/1\text{-}F_{ST}$ values) are plotted for each distance class against the logarithm of geographic distance between populations. Bars represent standard error at 95% level within each distance class

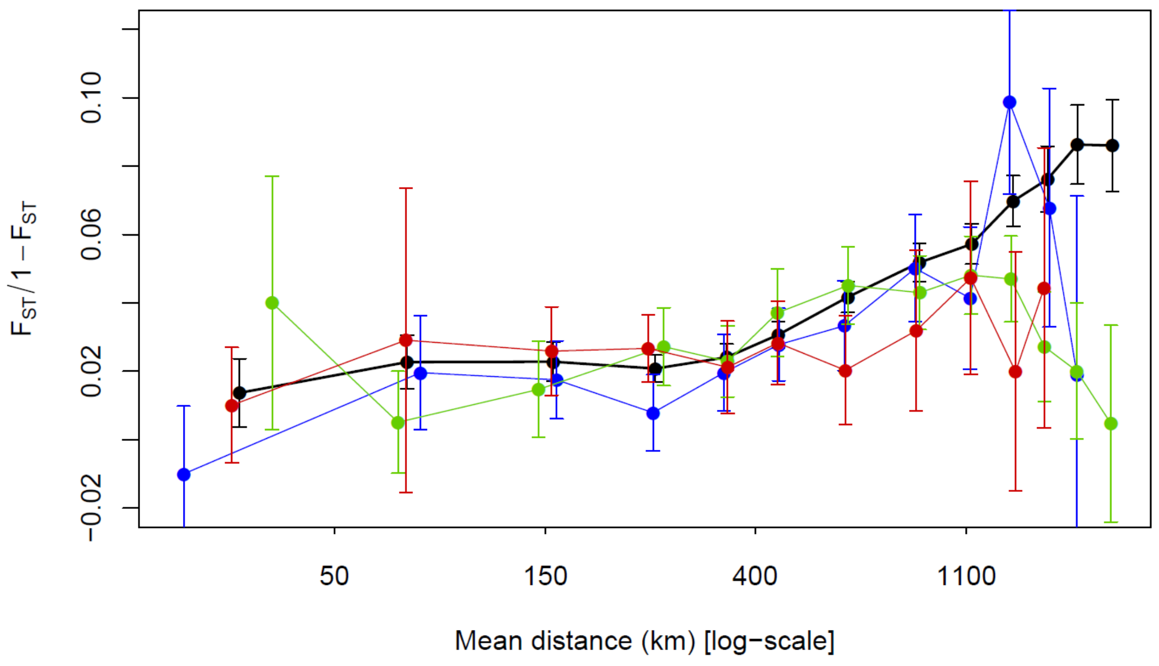

### Table S4: Outlier SNPs detected with a)PCAdapt and b)LEA.

**p-value**: p-value of PCAdapt/LEA test statistic. **pBonf**: p-value corrected for multiple comparisons using Bonferroni approach; **pBH**: p-value corrected for multiple comparisons using Benjamini-Hochberg approach; **q-value**: direct (unbiased) estimate of the False Discovery Rate (FDR) associated with pBH; **localFDR**: Empirical Bayesian posterior probability that the null is true conditional on the observed p-value; **nfp** : Expected number of false positives as based on q-value

We show only outlier SNPs (ie, those having q-value associated with an expected number of false positives, nfp, equal to zero).

***(a) pcadapt***

| SNP | p-value | pBonf | pBH | q-value | localFDR |
|---|---|---|---|---|---|
| QB_c13549-857 | 9.52 $10^{-5}$ | 0.0172 | 0.0172 | 0.0172 | 0.0762 |

***(b) lea***

| SNP | p-value | pBonf | pBH | q-value | localFDR |
|---|---|---|---|---|---|
| **154_1_251** | 0.000090 | 0.0243 | 0.0243 | 0.0243 | 0.0272 |
| **QB_c13549-857** | 0.000262 | 0.0708 | 0.0354 | 0.0354 | 0.0466 |
| **QB_c10512-206** | 0.000493 | 0.1330 | 0.0443 | 0.0443 | 0.0651 |
| **154_1_715** | 0.001005 | 0.2714 | 0.0603 | 0.0603 | 0.0956 |
| **154_1_845** | 0.001116 | 0.3015 | 0.0603 | 0.0603 | 0.1012 |

### Table S5: Outlier SNPs detected with *lfmm*.

**β:** lfmm test statistic for the correlation between environmental variable and allelic frequencies; **p-value**: p-value of the regression model. **q-value**: direct (unbiased) estimate of the False Discovery Rate (FDR) associated with p-value; **lFDR**: Empirical Bayesian posterior probability that the null is true conditional on the observed p-value; **CLIM**: synthetic climatic variable from PCA on climatic variables.

We show only outlier SNPs (ie, those having q-value associated with an expected number of false positives, nfp, equal to zero).

| SNP | β | p.value | q.value | lFDR | CLIM |
|---|---|---|---|---|---|
| QB_c10517-841 | 0.179 | 0.000 | 0.033 | 0.033 | Temp1 |
| 39_225 | -0.180 | 0.001 | 0.033 | 0.040 | Temp1 |
| QB_c15642-205 | -0.164 | 0.001 | 0.033 | 0.050 | Temp1 |
| 21_243 | -0.162 | 0.001 | 0.033 | 0.052 | Temp1 |
| 150_2_924 | 0.167 | 0.001 | 0.033 | 0.054 | Temp1 |
| SB_c15868-233 | 0.164 | 0.001 | 0.033 | 0.055 | Temp1 |
| 91_2_1441 | 0.155 | 0.003 | 0.065 | 0.081 | Temp1 |
| QB_c10512-206 | -0.176 | 0.000 | 0.004 | 0.006 | Temp2 |
| 150_2_924 | -0.199 | 0.000 | 0.004 | 0.008 | Temp2 |
| 21_243 | 0.188 | 0.000 | 0.004 | 0.008 | Temp2 |
| 52_1_246 | 0.185 | 0.000 | 0.009 | 0.015 | Temp2 |
| 154_2_371 | 0.175 | 0.000 | 0.015 | 0.023 | Temp2 |
| 154_1_845 | -0.148 | 0.001 | 0.018 | 0.029 | Temp2 |
| QB_c15913-902 | -0.162 | 0.001 | 0.023 | 0.038 | Temp2 |
| QB_c6167-1062 | 0.165 | 0.001 | 0.023 | 0.040 | Temp2 |
| 92_352 | 0.159 | 0.001 | 0.027 | 0.049 | Temp2 |
| QB_c15913-724 | 0.160 | 0.002 | 0.027 | 0.052 | Temp2 |
| ctrlfagus_c13215-830 | 0.156 | 0.002 | 0.027 | 0.054 | Temp2 |
| QB_c17017-1048 | 0.155 | 0.002 | 0.027 | 0.055 | Temp2 |
| 142_143 | 0.154 | 0.002 | 0.029 | 0.060 | Temp2 |
| 50_232 | -0.215 | 0.000 | 0.010 | 0.008 | Temp3 |
| 92_352 | -0.205 | 0.000 | 0.011 | 0.013 | Temp3 |
| QB_c13152-130 | -0.190 | 0.000 | 0.015 | 0.023 | Temp3 |
| 27_485 | -0.186 | 0.000 | 0.015 | 0.024 | Temp3 |
| 7_186 | 0.192 | 0.000 | 0.015 | 0.026 | Temp3 |
| QB_c10512-206 | 0.161 | 0.000 | 0.016 | 0.031 | Temp3 |
| SB_c5654-1048 | -0.169 | 0.001 | 0.017 | 0.035 | Temp3 |
| ctrlfagus_c15935-232 | 0.166 | 0.001 | 0.022 | 0.046 | Temp3 |
| QB_c13130-798 | -0.177 | 0.001 | 0.022 | 0.048 | Temp3 |
| 150_2_924 | 0.171 | 0.001 | 0.026 | 0.057 | Temp3 |
| 154_2_371 | 0.168 | 0.002 | 0.032 | 0.067 | Temp3 |
| 66_698 | 0.206 | 0.000 | 0.005 | 0.013 | Precip1 |
| QB_c13152-130 | 0.178 | 0.000 | 0.013 | 0.038 | Precip1 |
| 133_306 | -0.147 | 0.001 | 0.056 | 0.099 | Precip1 |

| | | | | | |
|---|---|---|---|---|---|
| **19_206** | -0.153 | 0.001 | 0.056 | 0.106 | Precip1 |
| **SB_c13339-608** | -0.193 | 0.000 | 0.029 | 0.021 | Precip2 |
| **SB_c968-192** | 0.080 | 0.001 | 0.053 | 0.034 | Precip2 |
| **SB_c968-1354** | -0.108 | 0.001 | 0.053 | 0.045 | Precip2 |
| **155_2_911** | -0.152 | 0.001 | 0.053 | 0.048 | Precip2 |
| **SB_c968-719** | -0.073 | 0.002 | 0.053 | 0.053 | Precip2 |
| **QB_c10517-414** | 0.147 | 0.002 | 0.053 | 0.053 | Precip2 |
| **SB_c968-935** | 0.075 | 0.002 | 0.053 | 0.059 | Precip2 |
| **QB_c13406-208** | -0.147 | 0.002 | 0.053 | 0.061 | Precip2 |
| **SB_c7640-125** | 0.153 | 0.002 | 0.053 | 0.061 | Precip2 |
| **129_685** | 0.250 | 0.000 | 0.000 | 0.000 | Precip3 |
| **21_243** | 0.229 | 0.000 | 0.000 | 0.000 | Precip3 |
| **QB_c10517-414** | -0.216 | 0.000 | 0.000 | 0.000 | Precip3 |
| **150_2_924** | -0.227 | 0.000 | 0.000 | 0.000 | Precip3 |
| **91_2_1441** | -0.212 | 0.000 | 0.000 | 0.002 | Precip3 |
| **148_1_1411** | -0.213 | 0.000 | 0.000 | 0.002 | Precip3 |
| **91_2_57** | 0.199 | 0.000 | 0.001 | 0.005 | Precip3 |
| **SB_c15868-233** | -0.197 | 0.000 | 0.001 | 0.005 | Precip3 |
| **154_2_371** | 0.198 | 0.000 | 0.001 | 0.006 | Precip3 |
| **68_277** | -0.185 | 0.000 | 0.002 | 0.011 | Precip3 |
| **92_352** | 0.166 | 0.001 | 0.010 | 0.033 | Precip3 |
| **SB_c13643-626** | -0.159 | 0.001 | 0.019 | 0.049 | Precip3 |
| **154_1_845** | -0.138 | 0.002 | 0.020 | 0.054 | Precip3 |
| **62_1_148** | -0.153 | 0.002 | 0.021 | 0.058 | Precip3 |
| **154_1_715** | -0.134 | 0.002 | 0.021 | 0.059 | Precip3 |
| **QB_c10512-206** | -0.127 | 0.003 | 0.025 | 0.069 | Precip3 |

### Table S6: Outlier SNPs detected with *Samβada*.

**G:** genotype**; D:** log-likelihood ratio test statistic deriving from the comparison among the null and alternative model for the *i-th* genotype**; $\beta_{clim}$:** Regression coefficient linking the climatic variable with the i-th genotype frequency; **p-value**: p-value associated with the log-likelihood test statistic D; **q-value**: direct (unbiased) estimate of the False Discovery Rate (FDR) associated with p-value; **lFDR**: Empirical Bayesian posterior probability that the null is true conditional on the observed p-value; **Clim**: synthetic climatic variable from PCA on climatic variables. We show only outlier SNPs (ie, those having q-value associated with an expected number of false positives, nfp, equal to zero).

| SNP | G | D | $\beta_{clim}$ | $\beta_{DAPC1}$ | $\beta_{DAPC2}$ | p.value | q.value | lFDR | OR_env | Clim |
|---|---|---|---|---|---|---|---|---|---|---|
| **154_1_251** | CC | 16.54 | 0.34 | -0.30 | -0.30 | 0.000 | 0.028 | 0.042 | 1.40 | Temp1 |
| **QB_c10460-202** | GG | 12.99 | -0.23 | 0.35 | 0.35 | 0.000 | 0.076 | 0.073 | 0.79 | Temp1 |
| **91_2_1441** | GT | 12.30 | 0.30 | -0.04 | -0.04 | 0.000 | 0.076 | 0.084 | 1.35 | Temp1 |
| **154_1_845** | TT | 11.90 | 0.33 | -0.46 | -0.46 | 0.001 | 0.076 | 0.092 | 1.39 | Temp1 |
| **154_1_390** | GG | 11.66 | -0.19 | 0.06 | 0.06 | 0.001 | 0.076 | 0.097 | 0.83 | Temp1 |
| **52_1_246** | TT | 22.04 | 0.28 | 0.10 | 0.10 | 0.000 | 0.002 | 0.005 | 1.32 | Temp2 |
| **ctrlfagus_c13215-830** | GG | 16.64 | 0.31 | 0.07 | 0.07 | 0.000 | 0.013 | 0.022 | 1.36 | Temp2 |
| **52_1_246** | AT | 16.44 | -0.24 | -0.06 | -0.06 | 0.000 | 0.013 | 0.024 | 0.79 | Temp2 |
| **150_2_924** | TT | 15.45 | 0.23 | -0.03 | -0.03 | 0.000 | 0.013 | 0.034 | 1.26 | Temp2 |
| **21_243** | TT | 15.26 | -0.24 | -0.02 | -0.02 | 0.000 | 0.013 | 0.036 | 0.79 | Temp2 |
| **154_2_371** | CC | 15.07 | 0.24 | 0.15 | 0.15 | 0.000 | 0.013 | 0.039 | 1.28 | Temp2 |
| **QB_c15913-902** | GG | 12.93 | -0.21 | -0.18 | -0.18 | 0.000 | 0.032 | 0.084 | 0.81 | Temp2 |
| **133_306** | CC | 12.82 | 0.33 | -0.44 | -0.44 | 0.000 | 0.032 | 0.088 | 1.39 | Temp2 |
| **133_306** | CT | 11.67 | -0.32 | 0.38 | 0.38 | 0.001 | 0.053 | 0.121 | 0.72 | Temp2 |
| **50_232** | AA | 22.70 | 0.47 | -0.24 | -0.24 | 0.000 | 0.001 | 0.005 | 1.60 | Temp3 |
| **50_232** | AG | 16.51 | -0.39 | 0.11 | 0.11 | 0.000 | 0.015 | 0.039 | 0.68 | Temp3 |
| **QB_c10512-206** | AA | 14.57 | -0.48 | -0.67 | -0.67 | 0.000 | 0.028 | 0.059 | 0.62 | Temp3 |
| **SB_c6451-300** | AA | 12.07 | -0.52 | -0.57 | -0.57 | 0.001 | 0.074 | 0.092 | 0.59 | Temp3 |
| **92_352** | TT | 11.82 | 0.33 | 0.15 | 0.15 | 0.001 | 0.074 | 0.096 | 1.40 | Temp3 |
| **QB_c7172-467** | TT | 14.46 | -0.25 | -0.02 | -0.02 | 0.000 | 0.088 | 0.075 | 0.78 | Precip1 |
| **133_306** | CC | 11.68 | -0.21 | -0.49 | -0.49 | 0.001 | 0.111 | 0.116 | 0.81 | Precip1 |
| **66_698** | CT | 11.43 | 0.31 | 0.02 | 0.02 | 0.001 | 0.111 | 0.123 | 1.36 | Precip1 |
| **66_698** | TT | 11.43 | -0.31 | -0.02 | -0.02 | 0.001 | 0.111 | 0.123 | 0.73 | Precip1 |
| **129_685** | AA | 22.81 | -0.53 | 0.08 | 0.08 | 0.000 | 0.001 | 0.002 | 0.59 | Precip3 |
| **21_243** | TT | 18.82 | -0.50 | 0.05 | 0.05 | 0.000 | 0.003 | 0.009 | 0.61 | Precip3 |
| **150_2_924** | TT | 16.35 | 0.43 | -0.09 | -0.09 | 0.000 | 0.008 | 0.018 | 1.54 | Precip3 |
| **SB_c13339-608** | CC | 14.13 | 0.62 | -0.01 | -0.01 | 0.000 | 0.017 | 0.028 | 1.85 | Precip3 |
| **SB_c13429-427** | AA | 14.00 | 0.60 | 0.07 | 0.07 | 0.000 | 0.017 | 0.028 | 1.82 | Precip3 |
| **91_2_1441** | TT | 12.62 | 0.69 | -0.07 | -0.07 | 0.000 | 0.030 | 0.036 | 1.99 | Precip3 |
| **91_2_1441** | GT | 11.80 | -0.67 | 0.06 | 0.06 | 0.001 | 0.040 | 0.043 | 0.51 | Precip3 |
| **134_2_834** | AA | 11.43 | -0.73 | 0.43 | 0.43 | 0.001 | 0.042 | 0.046 | 0.48 | Precip3 |
| **21_243** | CC | 11.24 | 0.53 | 0.07 | 0.07 | 0.001 | 0.042 | 0.049 | 1.70 | Precip3 |

### Table S7: Test for isolation by distance (IBD) and isolation by environment (IBE) based on variance partitioning and pRDA

For both sets of 218 putatively neutral SNPs (Neutral) and 52 outliers (Outliers), we first estimated the portions of variance (adjusted $R^2$) in the genetic structure explained by spatial ("Spat"), climatic ("Clim"), and joint spatio-climatic ("ClimSpat") structure (table A). We retained a variable number of principal coordinates in the PCoA analyses, explaining from 60 to 80% of total variance in genetic structure ("PCoA Explained Variance"). Significance of the variance components (Clim|Spat: of the climatic component when spatial component was accounted for; Spat|Clim: of the spatial component when climatic component was accounted for) was tested through ANOVA-like permutation test for partial redundancy analysis (pRDA), with result reported as F- and P-values.

Secondly, instead of a Climatic effect we considered distinct Temperature ("Temp") and Precipitation ("Prec") effects (Table B) for the set of 52 outliers and for the case where 80% of the total variance in genetic structure was explained. In this analysis,

**A- Temperature and Precipitation effects combined**

| Data | PCoA Explained Variance | Variance partitioning | | | | pRDA | | |
|---|---|---|---|---|---|---|---|---|
| | | Clim | ClimSpat | Spatial | Resid | Component | F-value | p-value |
| **Neutral** | 80% | 0.01 | 0.25 | 0.16 | 0.58 | Clim\|Spat | $F_{6,52}=1.1688$ | p=0.1641 |
| | | | | | | Spat\|Clim | $F_{5,52}=4.1193$ | p<0.0001 |
| **Outliers** | | 0.03 | 0.32 | 0.14 | 0.51 | Clim\|Spat | $F_{6,52}=1.6504$ | p=0.0058 |
| | | | | | | Spat\|Clim | $F_{5,52}=4.0076$ | p<0.0001 |
| **Neutral** | 70% | 0.02 | 0.28 | 0.18 | 0.53 | Clim\|Spat | $F_{6,52}=1.2933$ | p=0.0876 |
| | | | | | | Spat\|Clim | $F_{5,52}=4.8415$ | p<0.0001 |
| **Outliers** | | 0.04 | 0.36 | 0.15 | 0.46 | Clim\|Spat | $F_{6,52}=1.7775$ | p=0.0074 |
| | | | | | | Spat\|Clim | $F_{5,52}=4.794$ | p<0.0001 |
| **Neutral** | 60% | 0.01 | 0.33 | 0.18 | 0.47 | Clim\|Spat | $F_{6,52}=1.2728$ | p=0.1377 |
| | | | | | | Spat\|Clim | $F_{5,52}=5.3762$ | p<0.0001 |
| **Outliers** | | 0.03 | 0.41 | 0.17 | 0.39 | Clim\|Spat | $F_{6,52}=1.6672$ | p=0.0230 |
| | | | | | | Spat\|Clim | $F_{5,52}=5.8495$ | p<0.0001 |

B- **Distinct Temperature and Precipitation effects instead of a combined Climatic effect**

| **Data** | **PCoA Explained Variance** | **Variance partitioning** | | **pRDA** | | |
|---|---|---|---|---|---|---|
| | | | | Component | F-value | p-value |
| **Outliers** | **80%** | Temp | 0.01 | Temp\|Prec,Spat | $F_{3,57}$=7.9543 | p<0.0001 |
| | | Prec | 0.01 | Prec\|Temp,Spat | $F_{3,57}$=3.1206 | p=0.0003 |
| | | Spat | 0.14 | Spat\|Temp,Prec | $F_{5,55}$=5.364 | p<0.0001 |
| | | TempPrec | 0.01 | | NA | |
| | | TempSpat | 0.21 | | NA | |
| | | PrecSpat | 0.06 | | NA | |
| | | TempPrecSpat | 0.05 | | NA | |
| | | Residuals | 0.51 | | NA | |

NA: the interaction effects cannot be tested in pRDA.

### Figure S5: Respective contributions of climatic and spatial structure to the genetic structure.

These Venn diagram summarize the portions of variance (adjusted $R^2$) in the genetic structure explained by spatial and climatic effect for both sets of 218 putatively neutral SNPs (Neutral) and 52 outliers (Outliers). In left column, the effects of Temperature and Precipitation were considered together ("Climate" effect) while they were separated in right column. The intersections illustrate the contribution of joint effects, but note that this intersection is not equivalent to an interaction in an ANOVA for instance, and relates to the portion explained by effects which cannot be disentangled from each other.

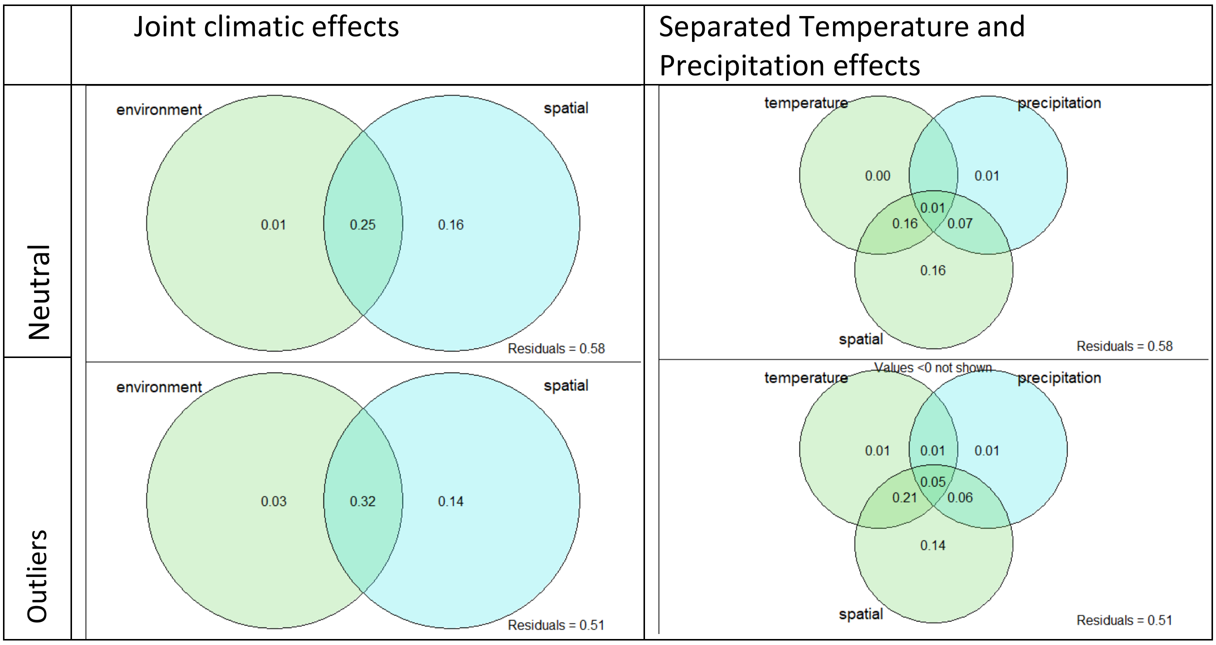

#### Table S8: Convergence with previous studies investigation genomic signature of selection at SNP loci.

For each previous study based on SNP markers, we give: **$N_{pop}$** = the number of studied populations/plots, with their location and sampling design. **$N_{CG}$**= the number of candidate genes with their origin (**Resource**) and the number of genotyped SNP (**$N_{SNP}$**). For each method (PD= population differentiation approach; EA= environmental association analyses; GPA= Genotype-Phenotype association), we list the number of outlier genes; the final column list the number of outlier genes in common with this study (CO).

| Study | $N_{pop}$ | Location | Sampling | $N_{CG}$ | Resource | $N_{SNP}$ | Method 1 | Outlier genes | Method 2 | Outlier genes | CO |
|---|---|---|---|---|---|---|---|---|---|---|---|
| Csilléry et al., 2014 | 4 | South-East France | Altitudinal gradient | 53 | Lalagüe *et al.*, 2014 | 546 | PD | 2-4 | Epistatic selection | 19 | 6 |
| Pluess et al., 2016 | 79 | Switzerland | Drought gradient | 52 | Lalagüe *et al.*, 2014 | 144 | EA | 19 | Logistic regression | 19 | 3 |
| Müller *et al.*, 2015b | 6 | Northern Germany | Translocation experiment | 9 | Müller *et al.*, 2015a | 46 | GPA | 6 | PD | 6 | 3 |
| Müller *et al.*, 2017 | 6 | Germany | Latitudinal gradient | 9 | Müller *et al.*, 2015a | 46 | GPA | 5 | | | 2 |
| Cuervo-Alarcon *et al.*, 2018 | 12 | Switzerland | Precipitation gradient | 24 | Lalagüe *et al.*, 2014; Seifert *et al.*, 2012 | 70 | PD | 12 | EA | 21 | 8 |
| Krajmerová *et al.*, 2017 | 19 | Slovenian glacial refugia | Provenance test | 6 | Lalagüe *et al.*, 2014; Seifert *et al.*, 2012 | 46 | PD | 3 | EA | 2 | 1 |
| Cuervo-Alarcon *et al.*, 2021 | 12 | Rhine and Rhône valleys in Switzerland | Precipitation gradients | 24 | Lalagüe et al., 2014; Seifert *et al.*, 2012 | 70 | GPA | 6 | | | |
