## Appendix 3 for "Genetic signatures of divergent selection in European beech (*Fagus sylvatica L.*) are associated with the variation in temperature and precipitation across its distribution range"

### Supplementary Material- Online Appendix 3: Principal Component Analyses of bioclimatic variables

This appendix details the results of Principal Component Analyses (PCA) performed on the set of 19 bioclimatic variables measured in the 64 populations (values given in Table S1). The objective was to obtain a limited number of ecological meaningful variables to include in gene-environment association analyses (GEA).

#### I. Data

At each sampling site, we extracted a set of 19 bioclimatic variables from the WorldClim database v.1.4 (Robert J. Hijmans, Cameron, Parra, Jones, & Jarvis, 2005) with a grid cell resolution of 30-arc second (ca 1×1 km) using DIVA-GIS v.7.5 (R J Hijmans, Guarino, Cruz, & Rojas, 2001). We first attempt to run PCA on the full set of bioclimatic variables, mixing those related to temperature (T in column “Type” of Table A2.1) and precipitations (P in column “Type” of Table A2.1)

| WordClim code | Variable (unit) | Type | Code |
|---|---|---|---|
| **BIO1** | Mean Annual Temperature (°C) | T | MAT |
| **BIO2** | Mean diurnal range (°C) | T | TDR |
| **BIO3** | Isothermality (BIO1/BIO7) * 100 | T | isoT |
| **BIO4** | Temperature Seasonality (CV) | T | Tseas |
| **BIO5** | Max Temperature of Warmest Period (°C) | T | MaxTWarm |
| **BIO6** | Min Temperature of Coldest Period (°C) | T | MaxTCold |
| **BIO7** | Temperature Annual Range (BIO5-BIO6; °C) | T | TAR |
| **BIO8** | Mean Temperature of Wettest Quarter (°C) | T | MeanTWet |
| **BIO9** | Mean Temperature of Driest Quarter (°C) | T | MeanTdry |
| **BIO10** | Mean Temperature of Warmest Quarter (°C) | T | MeanTwarm |
| **BIO11** | Mean Temperature of Coldest Quarter (°C) | T | MeanTcold |
| **BIO12** | Annual Precipitation (mm) | P | MAP |
| **BIO13** | Precipitation of Wettest Period (mm) | P | Pwet |
| **BIO14** | Precipitation of Driest Period (mm) | P | Pdry |
| **BIO15** | Precipitation Seasonality (CV; mm) | P | Pseas |
| **BIO16** | Precipitation of Wettest Quarter (mm) | P | PwetQ |
| **BIO17** | Precipitation of Driest Quarter (mm) | P | PdryQ |
| **BIO18** | Precipitation of Warmest Quarter (mm) | P | PwarmQ |
| **BIO19** | Precipitation of Coldest Quarter (mm) | P | PcoldQ |

**Table A2.1:** List of bioclimatic variables with their code (in Wordclim and in this study) and type (T: temperature-related; P: precipitation-related).

Spearman's rank correlations between all pairwise combinations of the 19 bioclimatic variables were calculated and visualized with the corrplot.mixed function from R package *corrplot* (Wei & Simko, 2017 ; Figure A2.1). Correlations occurred preferentially between temperature-related bioclimatic variables on the one hand and precipitation-related bioclimatic variables the other hand. By contrast, correlations between temperature-related versus precipitation-related bioclimatic variables were weaker.

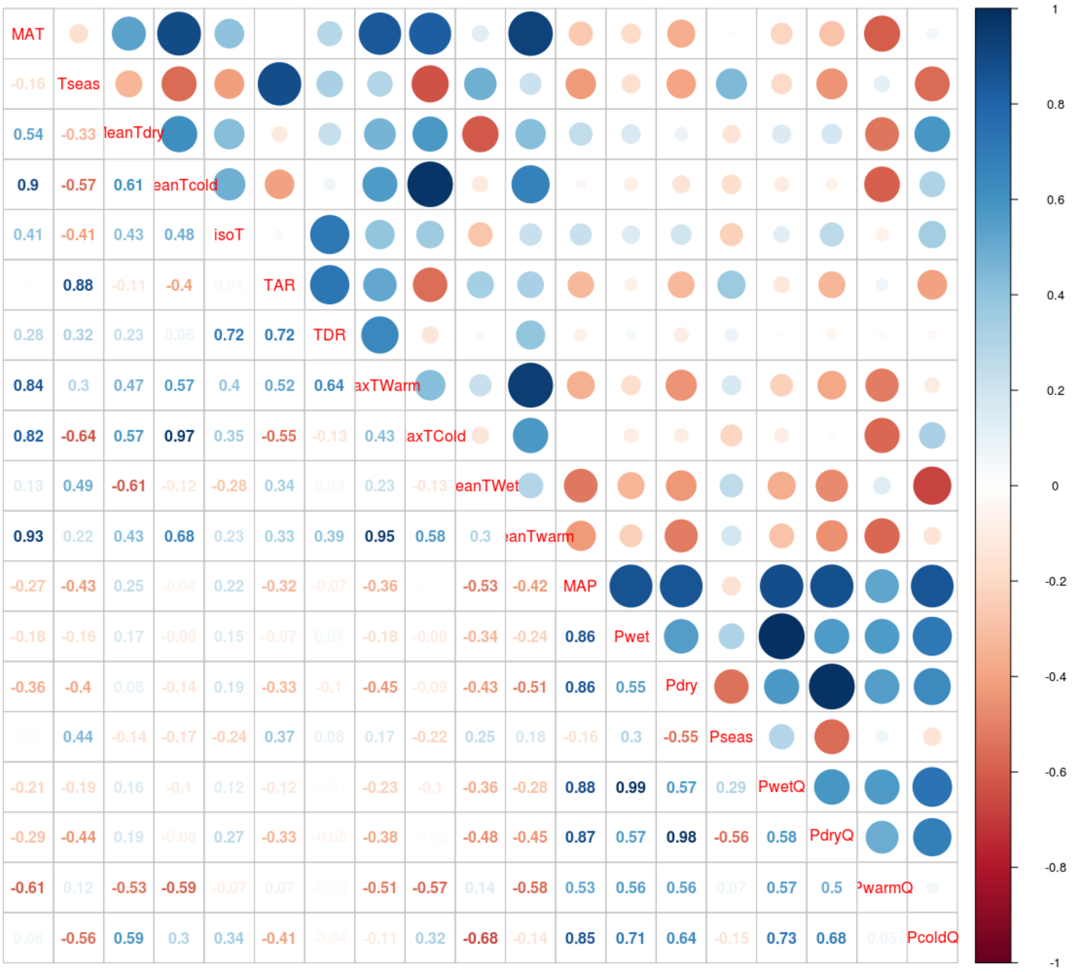


**Figure A2.1:** Pairwise correlations between all bioclimatic variables (see variable codes in Table A2.1)

#### II. PCA of the 11 bioclimatic variables related to temperature

##### 1. Raw correlations between variables

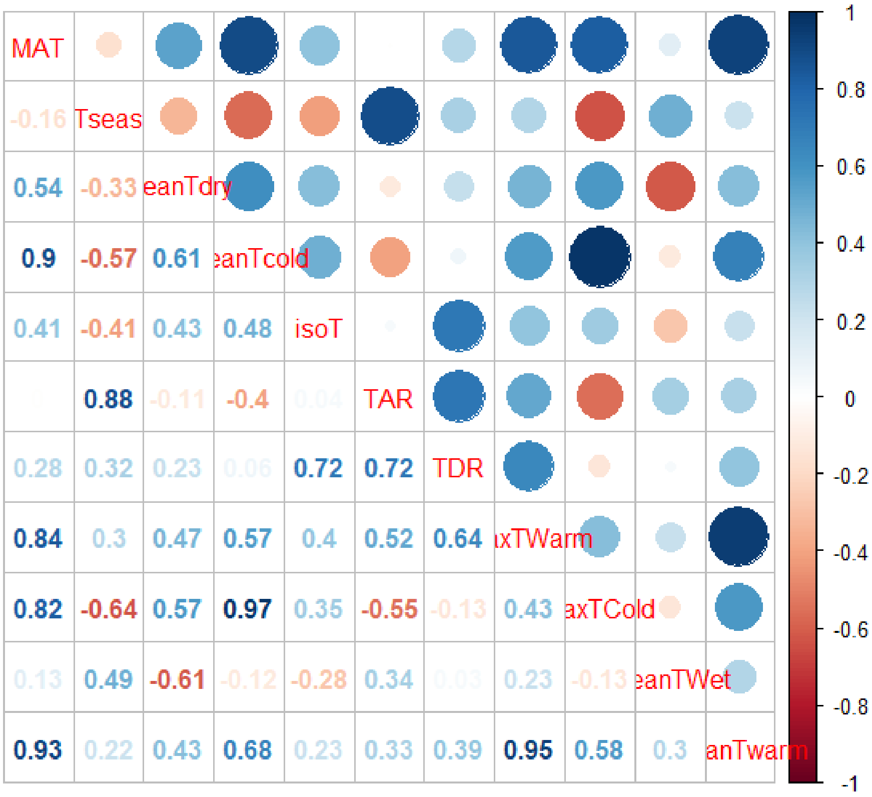


**Figure A2.5:** Pairwise correlations between temperature-related bioclimatic variables (see variable codes in Table A2.1)

We observed strong positive correlations between:

- MAT and meanTwarm (0.93), meanTcold (0.9), maxTwarm (0.84) and maxTCold (0.82).
- Tseas and TAR (0.88).
- meanTwarm & maxTwarm(0.95)
- meanTcold & maxTCold (0.97).

Moreover, Tseas and TAR were negatively correlated with MaxTCold (-0.64 and -0.55 respectively). Finally, MeanTwet and MeanTdry were negatively correlated (-0.61)

#### 2. Choice of the number k of principal components to keep

| PC | eigenvalue | % of variance | cumulative % of variance |
|---|---|---|---|
| **PC 1** | 4.936 | 44.869 | 44.869 |
| **PC 2** | 3.369 | 30.632 | 75.501 |
| **PC 3** | 1.689 | 15.358 | 90.858 |
| **PC 4** | 0.859 | 7.809 | 98.667 |
| **PC 5** | 0.114 | 1.035 | 99.703 |
| **PC 6** | 0.018 | 0.161 | 99.863 |
| **PC 7** | 0.012 | 0.109 | 99.972 |

**Table A2.3:** Eigenvalue (EV) associated to each PC from 1 to 7.

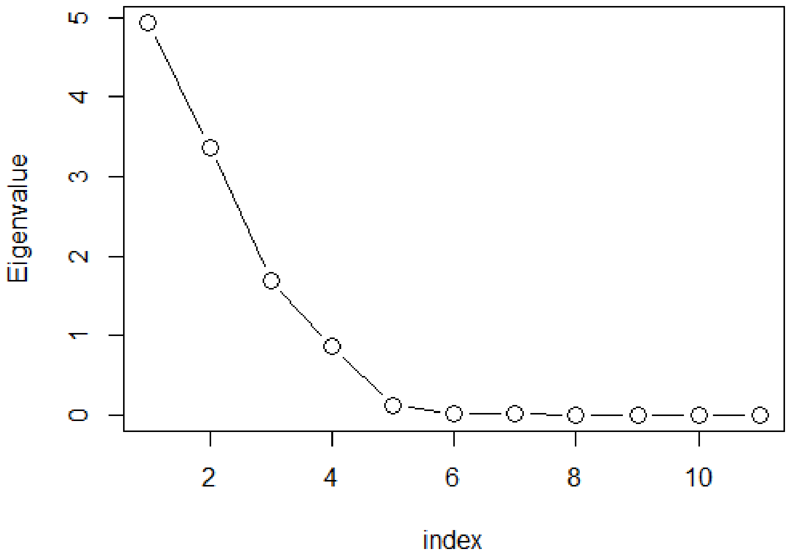


**Figure A2.6:** Ordered eigenvalues

The first four principal components (PC) captured 98.7% of the total variance (Table A2.3). The first three principal components account for 90.9% of the total variance of the dataset, and had eigenvalues >1. The rate of decrease of eigenvalues slows down after k=5 (Figure A2.6). Based on Kaiser-Guttman rule and considering the decrease in eigenvalues, we kept the first three PCs in this PCA on temperature-related variables, and renamed them Temp1, Temp2 and Temp3.

#### 3. Interpreting the PCA axes

(a)

(b)

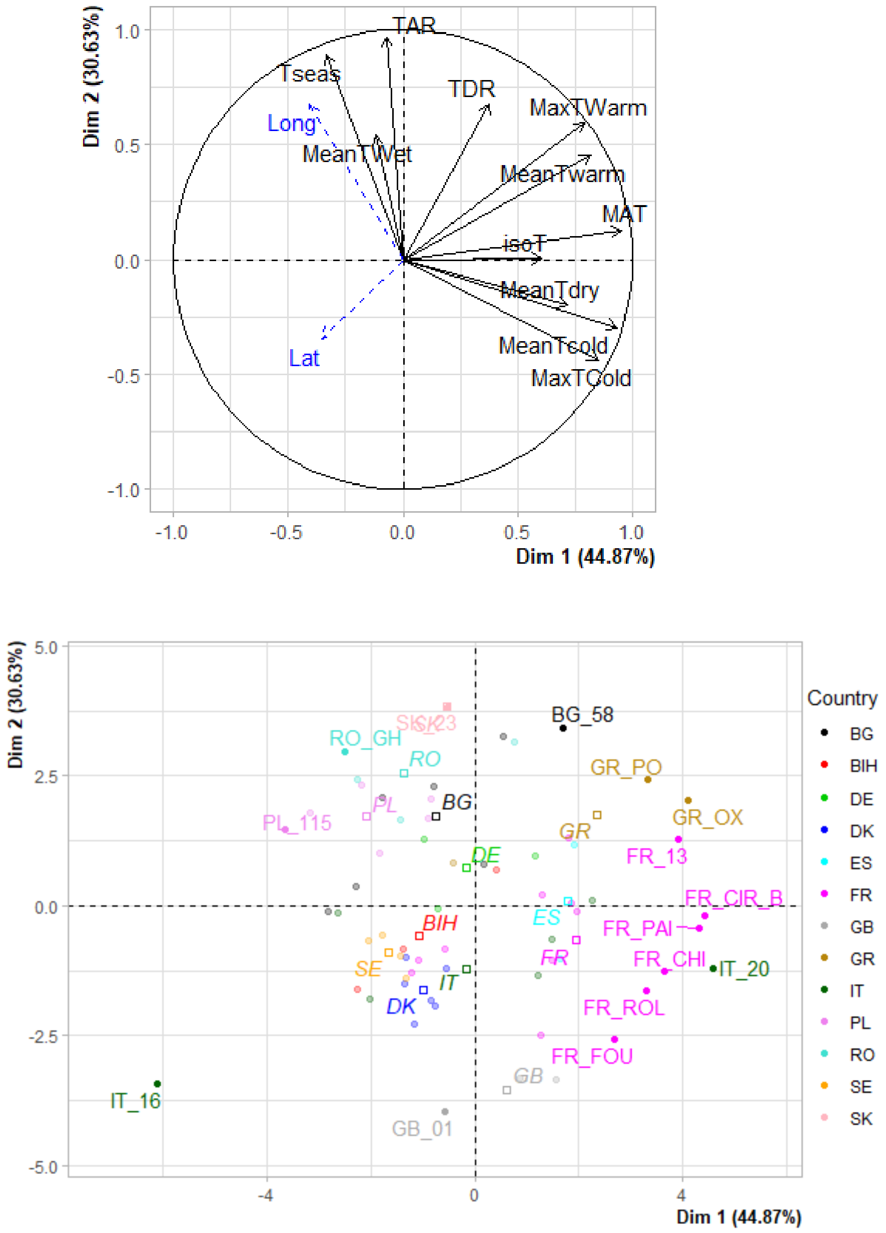


**Figure A2.7:** PCA graph of (a) variables and (b) populations for Temp1 and Temp 2. (a) Longitude and latitude were projected as supplementary variables. (b) Only the 15 populations that have the highest (squared) coordinates on Temp 1 and Temp 2 were drawn.

Temp1 is mostly driven by MAT and MeanTcold (Figure A2.7a). Temp1 is thus related to the variation of **mean temperatures**, opposing locations with hot (on the right) versus cold (on the left) climates. Extreme populations on Temp1 are located in Greece, France and central Italy at the right side and in Italy (IT-16), Poland (PL_115), Sweden and Denmark at the left side (Figure A2.7b).

Temp2 is mostly driven by TAR and Tseas, which are correlated. Temp2 is thus related to the variation of temperature variability, opposing locations experimenting a strong variation of temperatures among years and seasons (top side) to locations experimenting a low variation of temperatures (bottom side). Extreme populations on Temp2 are located in Great Britain and France at the bottom (oceanic climate), and Slovakia, Romania and Poland at the top (continental climate; Figure A2.7b). Longitude is well correlated with Temp2 (r=0.67). Temp2 can thus be interpreted as an axis of **climate continentality**.

(a)

(b)

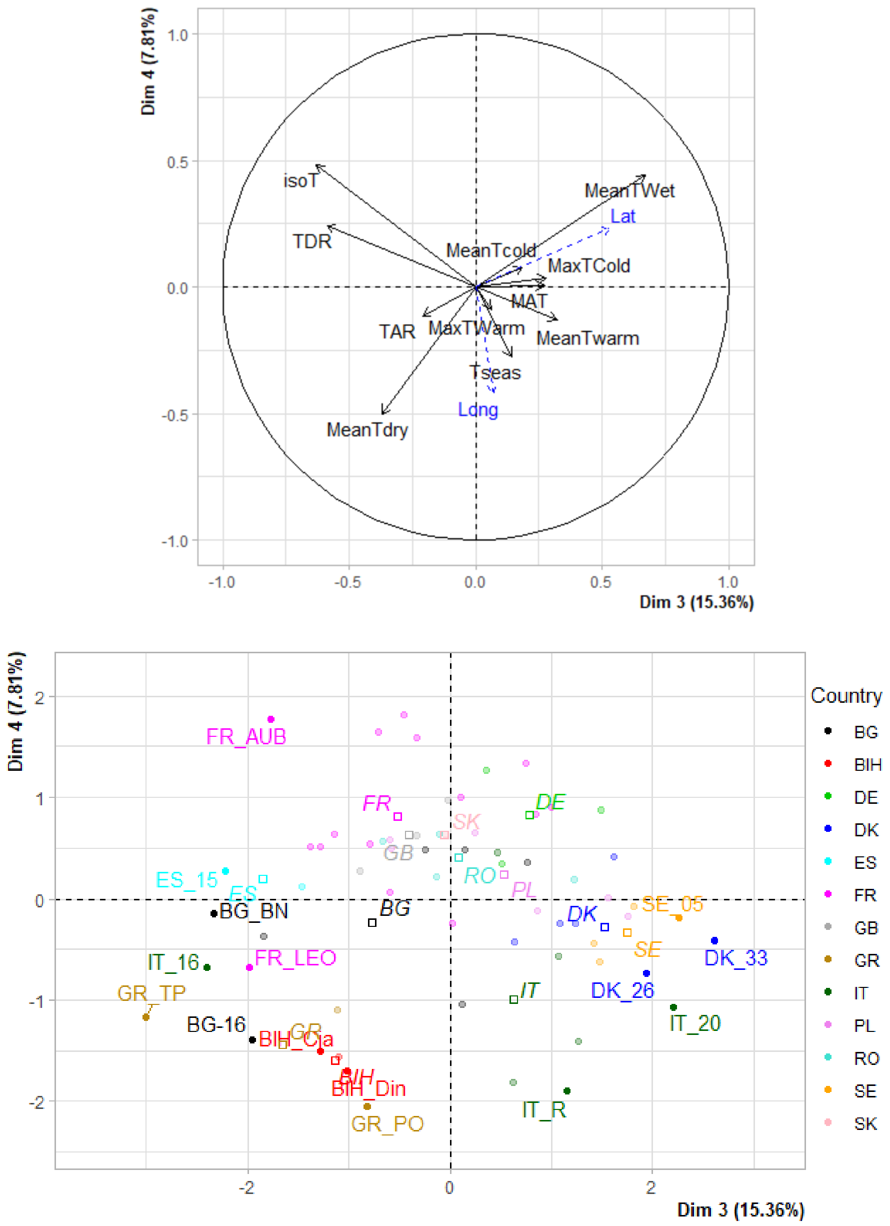


**Figure A2.8:** PCA graph of (a) variables and (b) populations for Temp3 and Temp4. (a) Longitude and latitude are projected as supplementary variables. (b) Only the 15 populations that have the highest (squared) coordinates on Temp3 and Temp4 are drawn.

Temp3 is driven by isoT, TDR and MeanTWet. Temp3 opposes locations with a high mean temperature during the wettest quarter of the year (MeanTWet) on the right, to locations with high isothermality and high temperature diurnal range on the left (Figure A2.8a). For those locations on the left, the wettest season corresponds to the coldest months (i.e., Mediterranean climates), and the diurnal range of temperatures is high as compared to the annual range of temperatures. By contrast, for locations on the right, the wettest season corresponds to the warmest months (i.e. temperate mesic climates) and the diurnal range of temperatures is low. Latitude is well correlated with Temp3 (r=0.53) Extreme populations on PC3 are located in Sweden and Denmark at the right side and in Italy, Greece, and France at the left side (Figure A2.8b). Temp3 can thus be interpreted as a **climate xericity axis,** opposing Mediterranean to temperate mesic climates.

#### III. PCA of the 8 bioclimatic variables related to precipitations

##### 1. Raw correlations between variables

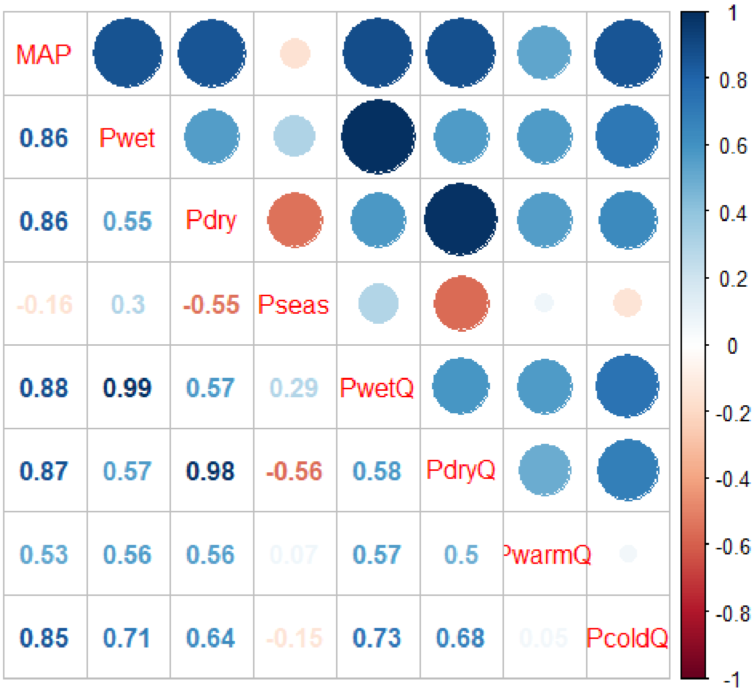


**Figure A2.9:** Pairwise correlations between precipitation-related bioclimatic variables (see variable codes in Table A2.1)

We observed strong positive correlations between:

- MAP and Pwet (0.86), Pdry (0.86), PwetQ (0.88), PdryQ (0.87) and PcoldQ (0.85).
- Pwet and PwetQ (0.99)
- Pdry and PdryQ (0.98).

##### 2. Choice of the number k of principal components to keep

| PC | eigenvalue | % of variance | cumulative % of variance |
|---|---|---|---|
| **PC 1** | 5.134 | 64.172 | 64.172 |
| **PC 2** | 1.789 | 22.365 | 86.537 |
| **PC 3** | 0.965 | 12.060 | 98.597 |
| **PC 4** | 0.047 | 0.585 | 99.182 |
| **PC 5** | 0.039 | 0.493 | 99.675 |

**Table A2.4:** Eigenvalue (EV) associated to each PC from 1 to 5.

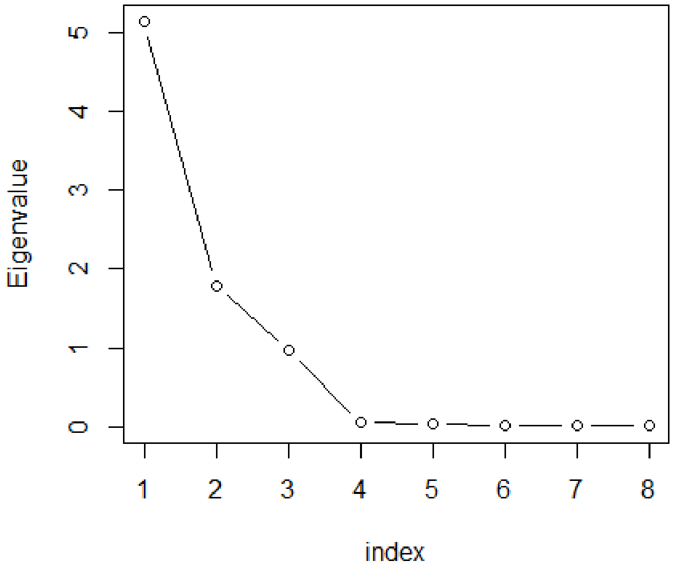


**Figure A2.10:** Ordered eigenvalues

The first four principal components (PC) captured 99.2% of the total variance (Table A2.4). The first three principal components account for 98.6% of the total variance of the dataset. The rate of decrease of eigenvalues slows down after k=3 (Figure A2.10). Based on the Kaiser-Guttman rule and considering the decrease in eigenvalues, we kept the first three PCs in this PCA on precipitation-related variables, and renamed them Precip1, Precip2, and Precip3.

#### 3. Interpreting the PCA axes

(a)

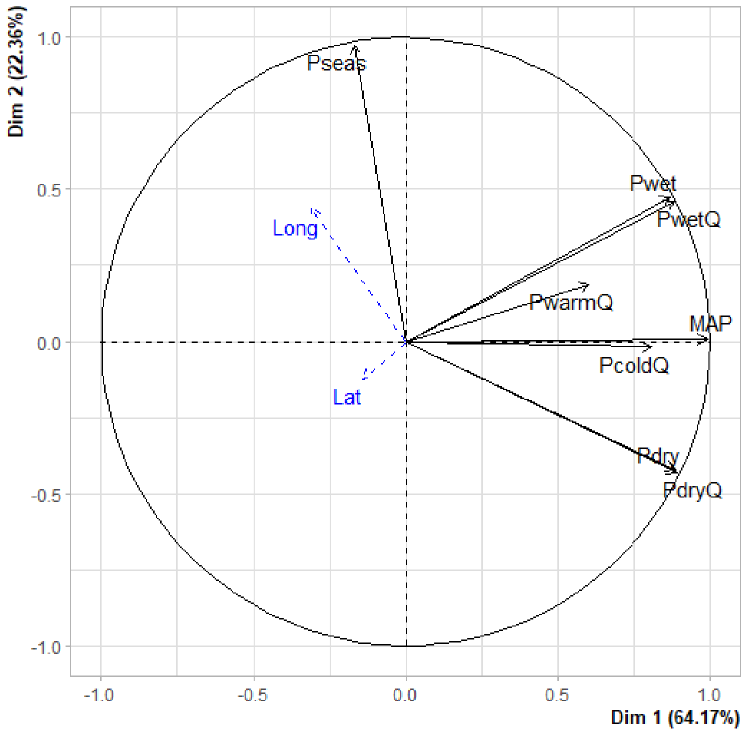

(b)

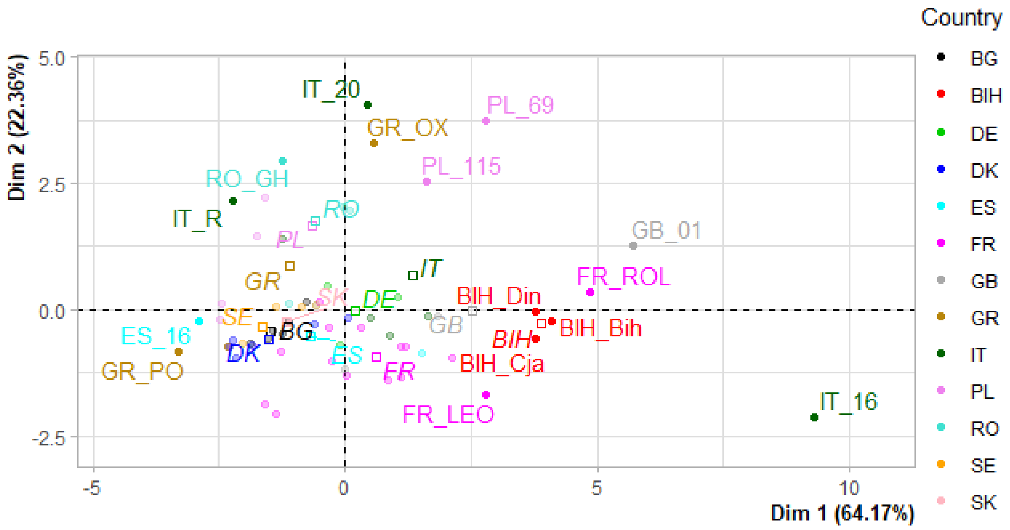


**Figure A2.11:** PCA graph of (a) variables and (b) populations for PC1 and PC2. (a) Longitude and latitude were projected as supplementary variables. (b) Only the 15 populations that have the highest (squared) coordinates on PC1 and PC2 were drawn.

Precip1 is mostly driven by MAP, and by the correlated variables Pdry, PwetQ, PdryQ (Figure A2.11a). Precip1 can be interpreted as a **precipitation abundancy axis**, opposing wet (on the right side) to dry (on the left side) climates. Extreme populations on Precip1 are located in Northern Italy (IT-16), Great Britain (GB_01) and Bosnia and Herzegovina at the right side and in Greece (GR_PO) and Spain (ES_16) at the left side (Figure A2.11b).

Precip2 is mostly driven by Pseas (Figure A2.11a). Precip2 can be interpreted as a **precipitation variability axis**, opposing climates with strong variability of precipitations among seasons (on the top side) to climate with low variability of precipitations (on the bottom side). Extreme populations on Precip2 are located in Italy (IT-20), Greece (GR_OX) and Poland at the top side and in France at the bottom side (Figure A2.11b).

(a)

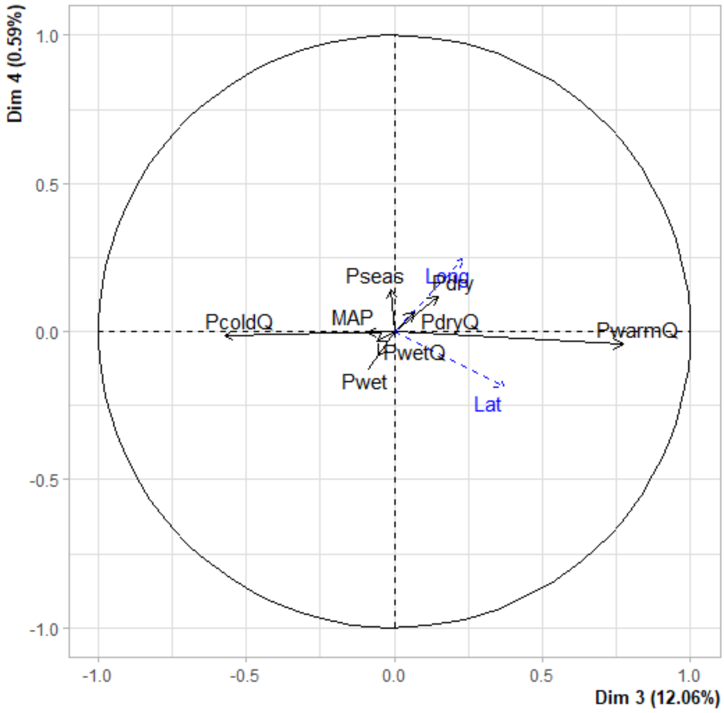


(b)

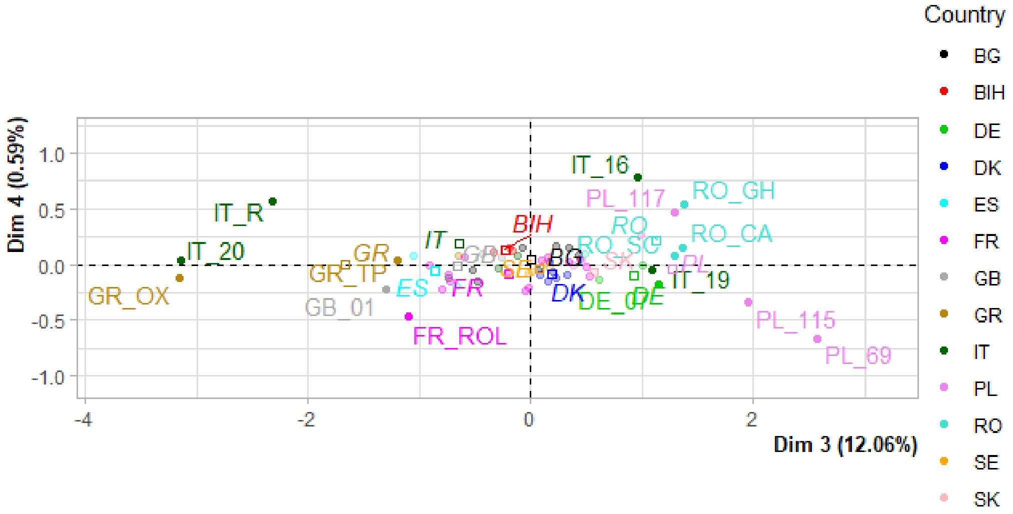


**Figure A2.12:** PCA graph of (a) variables and (b) populations for PRECIP3 and PRECIP4. (a) Longitude and latitude were projected as supplementary variables. (b) Only the 15 populations that have the highest (squared) coordinates on PRECIP3 and PRECIP4 were drawn.

Precip3 is mostly driven PwarmQ and PColdQ (Figure A2.12a). Precip3 hence captures the coupling between high precipitations and high seasonal temperatures, opposing climates where precipitations occur during the vegetation period (right side) to climates where precipitation occur rather in winter (left side). Extreme populations on Precip3 are located in Poland (PL_115, PL_69) and Romania (RO_GH, RO_CA) at the right side and in Greece (GR_OX) and Italy (IT_R, IT_20) at the left side (Figure A2.12b).

#### IV. Conclusions

We kept the three first PCs of the PCA on temperature-related variables (Temp1, Temp2, Temp3) and the three first PCs of the PCA on precipitations-related variables (Precip1, Precip2, Precip3). As expected, these 6 new synthetic variables are not correlated with each other within groups (ie, within temperature-related PC or within precipitations -related PCs), but there is a negative correlation between Temp1 and Precip3 (r=-0.66).

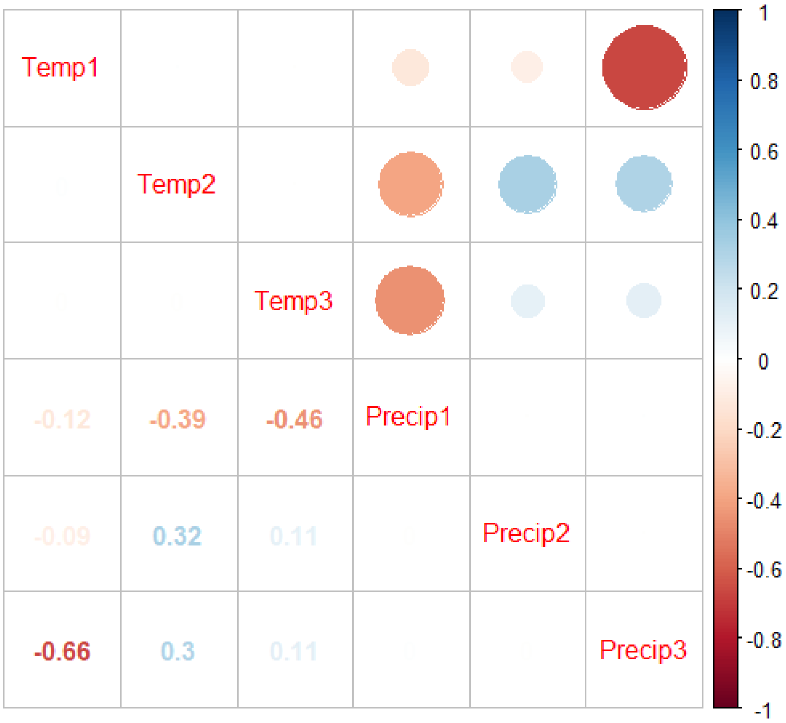
