## Appendix 2 for "Genetic signatures of divergent selection in European beech (*Fagus sylvatica L.*) are associated with the variation in temperature and precipitation across its distribution range"

This Appendix details the methods and parameters used to search for evidence of divergent selection. We used two different differentiation outlier approaches, aiming at identifying loci with unusual allele frequency differentiation among populations; and two different genotype-environment associations (GEA) approaches, aiming at identifying loci exhibiting significant correlations with ecological variables. For all test, we applied methods that explicitly model the effect of population structure, to remove eventual spurious associations arising from shared demographic history, patterns of isolation by distance, or cryptic relatedness.

### 1) Differentiation outlier analyses

#### a) pcadapt

The method implemented in the *R* package *pcadapt* (Luu, Bazin, & Blum, 2017) was used to test for association genetic loci and population structure and identify excessively associated loci (*i.e.* outliers bearing putative signatures of local adaptation). The software characterizes the genetic structure of populations through PCA, performs correlative models between individual allele counts and the most representative PCA dimension(s), estimates the strength of such an association through the robust Mahalanobis distance, and finally identifies outliers based on a chi-square distribution with degrees of freedom equal to the number of components retained.

The Cattell's rule was applied to identify a proper number of principal components to represent population structure.

The analysis was performed on the LD-pruned dataset in order to avoid the possible bias induced by linked loci during estimation of population structure.

#### b) lea

The second method we used takes advantage of the new $F_{ST}$ statistics developed by Martins et al. (2016) to identify outlier loci in admixed and in continuous populations, based on the computation of ancestry estimates obtained by the sparse non-negative matrix factorization (sNMF) algorithms implemented in the R package *LEA* v.2.0.0 (Frichot & François, 2015). Estimates of ancestry obtained by the sNMF algorithm can replace those obtained by the program structure advantageously for large SNP data sets.

The first step consists in selecting the most likely number of ancestral populations based on cross-entropy values for each K estimated by the *snmf()* function; here, we investigated K-values ranging from 1 to 8, with each K repeated 10 times.

The second step consists in computing the new $F_{ST}$ statistics using ancestry coefficients and ancestral allele frequencies as estimated by the *snmf()* function for the selected K value, and in converting them into absolute values of z-scores. The estimated z-scores require recalibration, based on the computation of the genomic inflation factor ($\lambda$) before applying a test for neutrality at each locus.

### 2) Genotype-Environment Associations (GEA) analyses

#### a) lfmm

The latent factor mixed model (LFMM) approach was used to test for associations between allele frequencies and environmental variables specified as fixed effects (Frichot, Schoville, Bouchard, & François, 2013). LFMM accounts for background population structure by incorporating relatedness as a random factor using unobserved variables (i.e., latent factors), which are equivalent to principal components of a PCA on allelic frequencies. LFMM has been shown to be an efficient and powerful method for different demographic scenarios, sampling designs, and in the case when loci are under weak selection (de Villemereuil, Frichot, Bazin, François, & Gaggiotti, 2014).

We used the most recent implementation of LFMM based on least squares estimates that was available in the R package *lfmm* (Caye & François, 2017). We estimated the LFMM parameters with the *lfmm_ridge()* function. The number of latent factors ($K_{lfmm}$) was obtained using the *snmf()* function of the *LEA* package as described above.

#### b) Samβada

The approach implemented by the software Samβada (v.0.8.0) was applied to scan the whole SNP dataset in search for significant genotype-environment associations (Stucki et al., 2017). Samβada relies on linear logistic regression to model the way genotypes are spread in the landscape as a function of habitat characteristics, and assumes loci displaying statistically significant association to underlie local adaptation. To identify such loci, log-likelihood ratio tests were devised where the performance of a null model including population structure only was compared with an alternative model including population structure plus one environmental variable at a time. *P*-values were then computed for each test from a chi-square distribution with one degree of freedom. Here, population structure was represented by the linear discriminant functions as estimated by DAPC.

### 3) Control of the false discovery rate

We followed the unified testing framework proposed by François, Martins, Caye, & Schoville (2016), which consists in recalibrating the test statistics by evaluating their expected value at selectively neutral loci; here, we used the local False Discovery Rate (FDR) method and determined, for each test, the *q*-value cutoff corresponding to zero expected false positives.

The FDR was controlled by translating *p*-values resulting from pcadapt, LEA, LFMM and Samβada analyses into *q*-values through the *R* package *qvalue* (Storey, Bass, Dabney, & Robinson, 2014/2020; Storey & Tibshirani, 2003). *Q*-value cut-offs were chosen so that no false positives were expected. In GEA, cut-offs were independently adapted for each environmental variable. The expected number of false positives associated with the i-*th* *q*-value ($FPD_i$) was calculated as $FPD_i=q_i \times n$, *n* being the number of tests with $q \leq q_i$. Only associations with *q*-value equal or lower than the zero expected false positives cut-off were deemed significant.

### 4) Assembly of outliers identified by different methods

We followed the strategy of Rellstab et al. (2015) to independently run the four methods and next to discuss the results by comparing lists of loci putatively under selection. Venn diagrams were produced with the Web-based tool InteractiVenn to visualize the consensus among the methods (Heberle, Meirelles, da Silva, Telles, & Minghim, 2015).
