## Appendix 1 for "Genetic signatures of divergent selection in European beech (*Fagus sylvatica L.*) are associated with the variation in temperature and precipitation across its distribution range"

### Raw dataset

We start from a raw dataset composed of 446 individuals:

```
nrow(data.table::fread("beechadapt.ped"))
```

```
## [1] 446
```

And 271 single-nucleotide polymorphisms (SNPs):

```
nrow(data.table::fread("beechadapt.map"))
```

```
## [1] 271
```

### Calculating missingness per individual and marker, and minor allele frequency

The sotware *plink* (Chang et al. 2015; Turner et al. 2011) is used to calculate missigness and minor allele frequency (MAF):

```
x <- "beechadapt" # Name of the dataset (without file extension)
thr <- seq(0, .3, .01) # Cut-offs
system(paste("./plink.exe --file ", x, " --missing --freq", sep=""))
```

```
## [1] 0
```

#### Individual call rate

The produced file *plink.imiss* contains missingness rate for each individual:

```
mind <- read.table("plink.imiss", stringsAsFactors = F, header = T)
knitr::kable(head(mind))
```

| FID | IID | MISS\_PHENO | N\_MISS | N\_GENO | F\_MISS |
| --- | --- | --- | --- | --- | --- |
| BG\_156 | BG\_156\_B | Y | 4 | 271 | 0.01476 |
| BG\_156 | BG\_156\_C | Y | 18 | 271 | 0.06642 |
| BG\_156 | BG\_156\_D | Y | 1 | 271 | 0.00369 |
| BG\_156 | BG\_156\_E | Y | 12 | 271 | 0.04428 |
| BG\_157 | BG\_157\_B | Y | 5 | 271 | 0.01845 |
| BG\_157 | BG\_157\_C | Y | 22 | 271 | 0.08118 |

Then, we can plot the expected number (and percentage) of removed individuals as a function of increasing missingness cut-offs:

```
y <- array(); y_ <- array()
for (i in 1:length(thr)) {
  y[i] <- (length(which(mind$F_MISS >= thr[i]))/nrow(mind))*100
  y_[i] <- length(which(mind$F_MISS >= thr[i]))
}; rm(i)
y <- as.data.frame(cbind(thr, y, y_))
colnames(y) <- c("Thr", "%_rm", "#_rm")
write.table(y, "mind_rm.txt", col.names = T, row.names = F, sep="\t", quote = FALSE)

plot(thr, y$`#_rm`, main = "",
     xlab = "Percentage of allowed missingness", ylab = "Removed individuals", 
     ylim = c(0, 100), t="l", las=1)
points(thr, y$`%_rm`, t="l", col="red")
abline(h=c(0, 10, 20), col="gray", lty=3)
abline(v=seq(0, .2, .05), col="gray", lty=3)
legend("topright", legend = c("Absolute number", "Percentage"), lty = 1, col=c("black", "red"), bg = "white")
```

We select a threshold of 15% missingness which allows both a sufficient amount of information to be retained for each individual and to remove not too much individuals (i.e., 16) due to low call rate:

```
mind_rm0.15 <- mind[which(mind$F_MISS>=0.15), ]
knitr::kable(mind_rm0.15)
```

|  | FID | IID | MISS\_PHENO | N\_MISS | N\_GENO | F\_MISS |
| --- | --- | --- | --- | --- | --- | --- |
| 11 | BG\_158 | BG\_158\_D | Y | 58 | 271 | 0.2140 |
| 25 | DE\_07 | DE\_07\_06 | Y | 51 | 271 | 0.1882 |
| 150 | GB\_03 | GB\_03\_08 | Y | 47 | 271 | 0.1734 |
| 158 | IT\_18 | IT\_18\_02 | Y | 50 | 271 | 0.1845 |
| 164 | IT\_18 | IT\_18\_10 | Y | 61 | 271 | 0.2251 |
| 240 | SE\_25 | SE\_25\_D | Y | 53 | 271 | 0.1956 |
| 251 | SK\_23 | SK\_23\_07 | Y | 231 | 271 | 0.8524 |
| 252 | SK\_23 | SK\_23\_08 | Y | 126 | 271 | 0.4649 |
| 269 | BG\_VT | BG\_VT\_10 | Y | 42 | 271 | 0.1550 |
| 284 | BIH\_Cja | BIH\_Cja\_6 | Y | 129 | 271 | 0.4760 |
| 302 | GR\_OX | GR\_OX\_8 | Y | 42 | 271 | 0.1550 |
| 303 | GR\_PO | GR\_PO\_5 | Y | 43 | 271 | 0.1587 |
| 311 | GR\_TP | GR\_TP\_5 | Y | 46 | 271 | 0.1697 |
| 315 | GR\_TP | GR\_TP\_9 | Y | 133 | 271 | 0.4908 |
| 318 | GR\_TP | GR\_TP\_12 | Y | 67 | 271 | 0.2472 |
| 334 | RO\_DB | RO\_DB\_8 | Y | 49 | 271 | 0.1808 |

```
write.table(mind_rm0.15, "mind_rm0.15.txt", col.names = T, row.names = F, 
            sep="\t", quote = FALSE)
mind_rm0.15 <- table(mind_rm0.15$FID)
```

We also inspect the impact of pruning in terms of number of individuals removed per population:

```
n.ind.pop <- data.table::fread("beechadapt.ped", header = F)
n.ind.pop <- table(n.ind.pop$V1)
n.ind.pop <- n.ind.pop[match(names(mind_rm0.15), 
                             names(n.ind.pop))]
n.ind.pop <- cbind(n.ind.pop, mind_rm0.15)
colnames(n.ind.pop) <- c("N.er of individuals per populations", "N.er of removed individuals") 
knitr::kable(n.ind.pop)
```

|  | N.er of individuals per populations | N.er of removed individuals |
| --- | --- | --- |
| BG\_158 | 4 | 1 |
| BG\_VT | 8 | 1 |
| BIH\_Cja | 8 | 1 |
| DE\_07 | 10 | 1 |
| GB\_03 | 10 | 1 |
| GR\_OX | 8 | 1 |
| GR\_PO | 8 | 1 |
| GR\_TP | 8 | 3 |
| IT\_18 | 8 | 2 |
| RO\_DB | 8 | 1 |
| SE\_25 | 4 | 1 |
| SK\_23 | 9 | 2 |

Removal seems to be spread among populations which remain well represented even after pruning.

#### SNP call rate

The produced file *plink.lmiss* contains missingness rate for each locus:

```
geno <- read.table("plink.lmiss", stringsAsFactors = F, header = T)
knitr::kable(head(geno))
```

| CHR | SNP | N\_MISS | N\_GENO | F\_MISS |
| --- | --- | --- | --- | --- |
| 1 | QB\_c10512206 | 15 | 446 | 0.03363 |
| 1 | QB\_c10517302 | 13 | 446 | 0.02915 |
| 1 | QB\_c10517414 | 10 | 446 | 0.02242 |
| 1 | QB\_c7172467 | 11 | 446 | 0.02466 |
| 1 | QB\_c7172995 | 11 | 446 | 0.02466 |
| 1 | QB\_c13086298 | 12 | 446 | 0.02691 |

Analogously to what done for individual missingness, we can plot the expected number (and percentage) of removed SNPs as a function of increasing missingness cut-offs:

```
y <- array(); y_ <- array()
for (i in 1:length(thr)) {
  y[i] <- (length(which(geno$F_MISS >= thr[i]))/nrow(geno))*100
  y_[i] <- length(which(geno$F_MISS >= thr[i]))
}; rm(i)
y <- as.data.frame(cbind(thr, y, y_))
colnames(y) <- c("Thr", "%_rm", "#_rm")
write.table(y, "geno_rm.txt", col.names = T, row.names = F, sep="\t", quote = FALSE)

plot(thr, y$`#_rm`, main = "",
     xlab = "Percentage of allowed missingness", ylab = "Removed SNPs", 
     ylim = c(0, 100), t="l", las=1)
points(thr, y$`%_rm`, t="l", col="red")
abline(h=c(0, 10, 20), col="gray", lty=3)
abline(v=seq(0, .2, .05), col="gray", lty=3)
legend("topright", legend = c("Absolute number", "Percentage"), lty = 1, col=c("black", "red"), bg = "white")
```

Again, a 15% missingness cut-off grants a sufficient amount of information to be retained and to discard one SNP only from the dataset due to low call rate:

```
geno_rm0.15 <- geno[which(geno$F_MISS>=0.15), ]
print(geno_rm0.15)
```

```
##    CHR          SNP N_MISS N_GENO F_MISS
## 98   1 QB_c13130173     99    446  0.222
```

```
write.table(geno_rm0.15, "geno_rm0.15.txt", col.names = T, row.names = F, 
            sep="\t", quote = FALSE)
```

#### Minor allele frequency

The produced file *plink.frq* contains MAF values for each locus:

```
maf <- read.table("plink.frq", stringsAsFactors = F, header = T)
knitr::kable(head(maf))
```

| CHR | SNP | A1 | A2 | MAF | NCHROBS |
| --- | --- | --- | --- | --- | --- |
| 1 | QB\_c10512206 | A | T | 0.3689 | 862 |
| 1 | QB\_c10517302 | A | G | 0.4642 | 866 |
| 1 | QB\_c10517414 | T | A | 0.1433 | 872 |
| 1 | QB\_c7172467 | T | G | 0.4724 | 870 |
| 1 | QB\_c7172995 | A | G | 0.3713 | 870 |
| 1 | QB\_c13086298 | A | C | 0.2523 | 868 |

Then, we can plot the expected number (and percentage) of removed loci as a function of increasing MAF cut-offs:

```
y <- array(); y_ <- array()
for (i in 1:length(thr)) {
  y[i] <- (length(which(maf$MAF <= thr[i]))/nrow(geno))*100
  y_[i] <- length(which(maf$MAF <= thr[i]))
}; rm(i)
y <- as.data.frame(cbind(thr, y, y_))
colnames(y) <- c("maf", "%_rm", "#_rm")
write.table(y, "maf_rm.txt", col.names = T, row.names = F, sep="\t", quote = FALSE)

plot(thr, y$`#_rm`, main = "",
     xlab = "Allowed MAF", ylab = "Removed SNPs", 
     xlim = c(0, 0.2), ylim = c(0, 100), t="l", las=1)
points(thr, y$`%_rm`, t="l", col="red")
abline(h=c(0, 10, 20), col="gray", lty=3)
abline(v=seq(0, .2, .05), col="gray", lty=3)
legend("topleft", legend = c("Absolute number", "Percentage"), lty = 1, col=c("black", "red"),
       bg="white")
```

### Dataset pruning

In order to prune the dataset, we specify the selected QC cut-off for individual and SNP missingness as follow:

```
mind <- .15 # allowed missingness for individuals
geno <- .15 # allowed missingness for SNPs
```

A standard MAF cut-off of 1% leads to remove no SNP (see fig. above), so that we can actually avoid including such a step in subsequent analyses. Then, we can use *plink* to remove individuals and loci with call rate lower than the selected threshold (85% in both cases):

```
# .ped/.map format
system(paste("./plink.exe --file ", x,
             " --mind ", mind,
             " --geno ", geno,
             " --recode --out ", paste(x, "_mind", mind, "geno", geno, sep=""), 
             sep=""))
```

```
## [1] 0
```

```
# plink binary format (.bed/.bim/.fam)
system(paste("./plink.exe --file ", 
             paste(x, "_mind", mind, "geno", geno, sep=""),
             " --make-bed --out ", paste(x, "_mind", mind, "geno", geno, sep=""), 
             sep=""))
```

```
## [1] 0
```

We can then compute the number of individuals and loci in the QC-ed dataset:

```
nrow(data.table::fread("beechadapt_mind0.15geno0.15.fam"))
```

```
## [1] 430
```

```
nrow(data.table::fread("beechadapt_mind0.15geno0.15.bim"))
```

```
## [1] 270
```

Further, we can inspect populations’ consistency in the pruned dataset (i.e., the number of individuals per population):

```
n.ind.pop <- sort(table(data.table::fread("beechadapt_mind0.15geno0.15.fam")$V1))
par(mfrow=c(2, 1), oma = c(1, 2, 1, 2), mar = c(4, 4, 0.5, 0.5))
barplot(n.ind.pop, names.arg = names(n.ind.pop), col = "gray", 
        ylab = "# ind.s/pop.", border = T, las = 2, cex.names = 0.4)
hist(n.ind.pop, main = "", xlab = "Frequency classes", col = "gray", las = 1)
```

The mean number of individuals per population in the QC-ed datset is:

```
round(mean(n.ind.pop))
```

```
## [1] 7
```

The standard deviation:

```
round(sd(n.ind.pop))
```

```
## [1] 2
```

The ranges in population consistency (min/max number of individuals per population) are:

```
range(n.ind.pop)
```

```
## [1]  3 10
```

### Conclusion

The QC-ed dataset comprises 430 individuals and 270 SNPs. On average, we have 7 (±2) individuals per population; the less represented populations are composed by 3 individuals, the most represented ones by 10 individuals.
